# Structural basis of the π-stacking network governing cofactor-substrate cooperativity of SbSOMT

**DOI:** 10.1101/2025.10.22.683835

**Authors:** Kah Chee Pow, Nan Zhang, Ming Yan, Xiaorong Wang, Andy C. W. Lui, Clive Lo, Quan Hao

**Affiliations:** Spallation Neutron Source Science Center, Dongguan, Guangdong, China; Institute of High Energy Physics, Chinese Academy of Sciences, Beijing, China; School of Biomedical Sciences, LKS Faculty of Medicine, The University of Hong Kong, Pokfulam, Hong Kong, China; Proteomics and Metabolomics Facility, Cornell University, Ithaca, NY, USA; School of Biological Sciences, The University of Hong Kong, Pokfulam, Hong Kong, China

**Author notes:** These authors contributed equally to this work (Kah Chee Pow, Nan Zhang).

**Keywords:** cooperativity, binding kinetics, methyltransferase, π-stacking network, structural dynamics, X-ray crystallography

## Abstract

SAM-dependent methyltransferases are ubiquitous enzyme catalyzing methylation of substrate by consuming *S*-adenosyl methionine (SAM) cofactor. In this study, we discovered the positive cofactor-substrate cooperativity of SbSOMT and uncovered the underlying mechanism at structural basis. The binding kinetics analyses show that SbSOMT exhibits bilateral positive cooperativity except the unproductive pairing between SAM and product due to the steric hindrance of an additional methyl group. Unprecedentedly, two cofactor-analogous inhibitors, *S*-adenosyl homocysteine (SAH) and sinefungin demonstrated opposite effects on substrate binding kinetics while attaining positive cooperativity. SAM-resembling sinefungin demonstrated higher foldchange in the acceleration of substrate association rate constant than deceleration of dissociation rate constant. Integrating SbSOMT structural insights at multiple states, we identified the dynamical W279-driven π-stacking network governs the cofactor-substrate cooperativity. SbSOMT exhibits closed conformation at first-ligand bound states, where the C-terminal W279 serves as the central plane for π-π interaction between N-terminal H196, catalytic H282, and/or substrate. We propose that the binding of first ligand overcome the energy barrier represented by W279 π-stacking network, leading to the conformational landscapes in favor to second binding event. Our study highlights the intrinsic design of SbSOMT in pertaining enzyme competency and serves as a foundation to expand the knowledge and applications of cofactor-substrate cooperativity of the structurally diverse methyltransferases.

## Introduction

SAM-dependent methyltransferases are ubiquitous enzyme that recruits *S*-adenosyl methionine (SAM), as the methyl-donor cofactor for substrate methylation. Cofactor-substrate cooperativity describes the interplay of methyltransferase, cofactor, and substrate. Proteins are intrinsically dynamic, existing in conformational ensembles with switchable conformational landscapes upon ligand(s) binding. Cooperativity arises when the binding of the first ligand leads to a second binding event with perturbed binding kinetics.

Among methyltransferases, applications of cooperativity are thriving ahead of fundamental studies concerning the underlying mechanism. Emerging studies have strategically manifested cofactor-dependent cooperativity to amplify affinity and selectivity of the inhibitors based on the differential metabolic flux in particular microenvironments (*1-5*). On the other hand, methyltransferases are attractive biocatalyst for late-stage modification of fine chemical; owning to the diversity and amenability from the aspects of substrate spectrum, chemo-selectivity, cofactor expandability and more (*6, 7*). SAM-dependent methyltransferases comprise diverse SAM binding domain architecture with the vast majority belonging to the Class I SAM-dependent methyltransferase superfamily (*8, 9*). The SAM binding domain of Class I SAM-dependent methyltransferase superfamily conserves a seven-β-strand core Rossmann fold with shared ancestry along several nucleoside binding domains that recognize nucleoside cofactors such as ATP, NAD, and GDP (*9, 10*). One Class I subgroup in particular, namely plant *O*-methyltransferases, has emerged to catalyze interesting chemical reactions such as pericyclic reaction, redox activity, and diazo installation (*11-15*).

Our previous study has uncovered the structural basis of the substrate regioselectivity in *Sorghum bicolor* stilbene *O*-methyltransferase (SbSOMT) and showed implications among plant *O*-methyltransferases (*16*). In this study, we ponder deeper into the structure-function relationship of cofactor-substrate cooperativity in SbSOMT. We combined binding kinetics analysis and X-ray crystallography to annotate the positive cofactor-substrate cooperativity with unprecedented binding kinetics properties, and provide a structural basis to address the underlying mechanism.

## Results

### SbSOMT exhibits bilateral positive cofactor-substrate cooperativity

To decipher the cofactor-substrate cooperativity, we employed isothermal titration calorimetry (ITC) to profile the binding affinity of substrate and cofactor analogs. Cooperativity (σ) can be calculated based on the fold-change of ligand binding affinity in the context of first binding event (apo) over ligand affinity in the context of second binding event (cognate-ligand-saturated); where cofactor is the cognate ligand of substrate and vice versa. Note that, SbSOMT catalytic cycle (fig. S1A) involves methylation of resveratrol (RSV) sequentially to produce mono-methylated pinostilbene (PNS) and di-methylated pterostilbene (PTR), while converting SAM to *S*-adenosyl homocysteine (SAH) per transfer (*16*). The three substrate analogs (RSV, PNS and PTR) and the two cofactor analogs (SAM and SAH) were probed in addition to sinefungin (SFG). SFG consists of a non-reactive amine group equivalent to the transferable methyl group of SAM (fig. S1B). Hence, SFG served as SAM analog to pair with reactive RSV for inferring the SAM-dependent characteristics distinct to the SAH-dependent characteristics.

The affinity profile (Table 1 and Fig. 1A) towards RSV has indeed shown distinct features between SFG-saturating and SAH-saturating conditions. In SAH-saturating condition, the titration was monophasic and fitted with affinity (K_D_) of 197 nM, showing a positive cooperativity of 12.03. Conversely, the thermogram of SFG-saturating condition (Fig. 1B) was showcasing biphasic binding kinetics. The titration curve was successfully fitted with a “two sets of sites” fitting model and the stoichiometry (N) of both sites were equally fitted at 0.39. The shift of stoichiometry and the biphasic thermogram imply that SFG-bound SbSOMT was dimeric and the protomers’ binding site has independent and different binding kinetics. The two sites demonstrate positive cooperativity at different scales and opposite binding enthalpy. At one protomer, the binding was endothermic (2.63 kcal/mol) with RSV affinity strengthened by ∼3-fold to 818 nM. The opposite exothermic binding (-5.48 kcal/mol) was fitted with a K_D_ of 24.2 nM and σ of 97.93; attaining the strongest affinity and cooperativity among all. The profile implies that the cofactor-bound SbSOMT exhibits positive cooperativity towards RSV in general, while SAM-bound SbSOMT positive cooperativity likely exhibits dimer asymmetry characteristic in resemblance to SFG-bound SbSOMT.

**Table 1.**
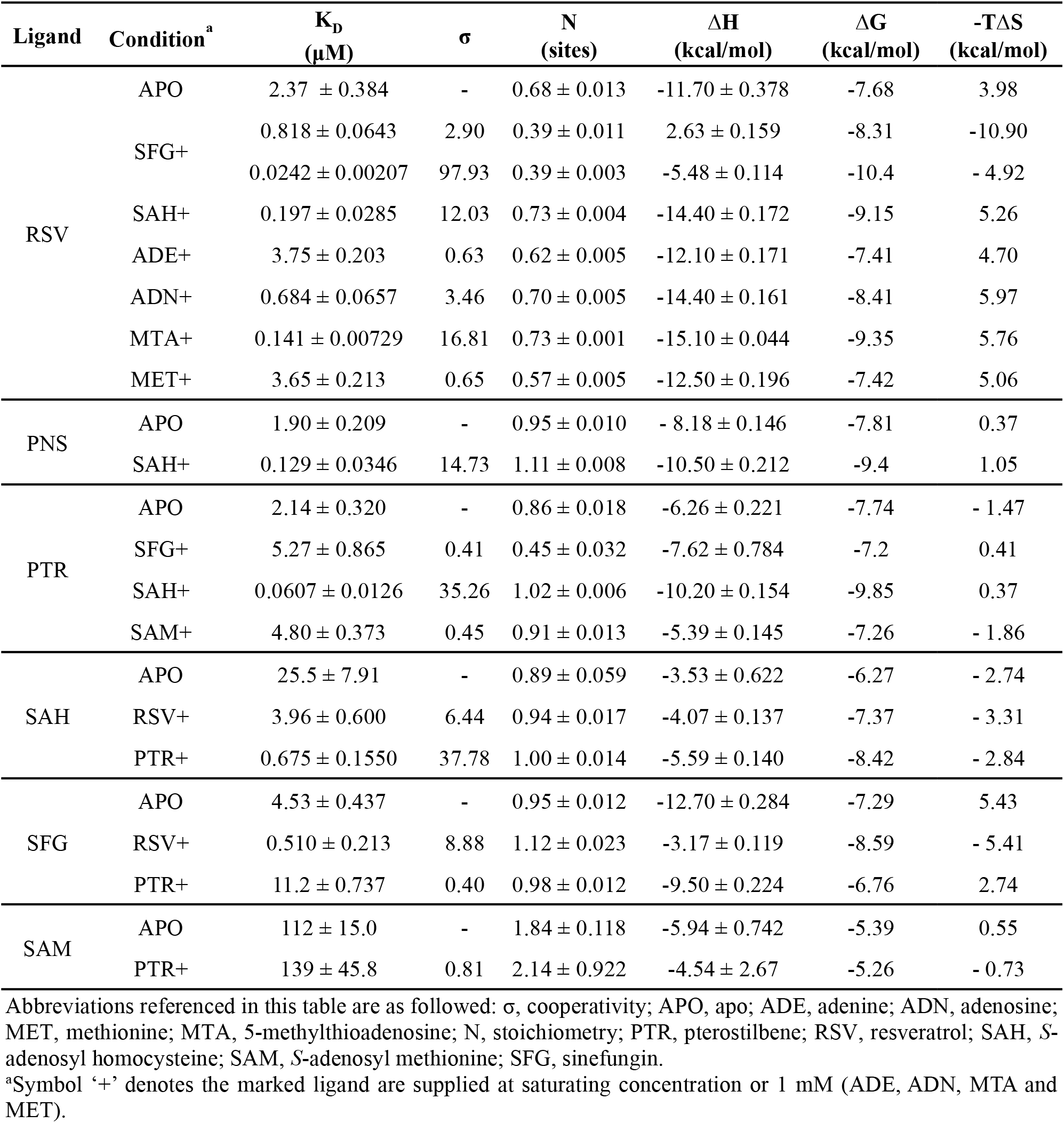
Binding affinity profile of SbSOMT measured by ITC.

**Fig. 1.**
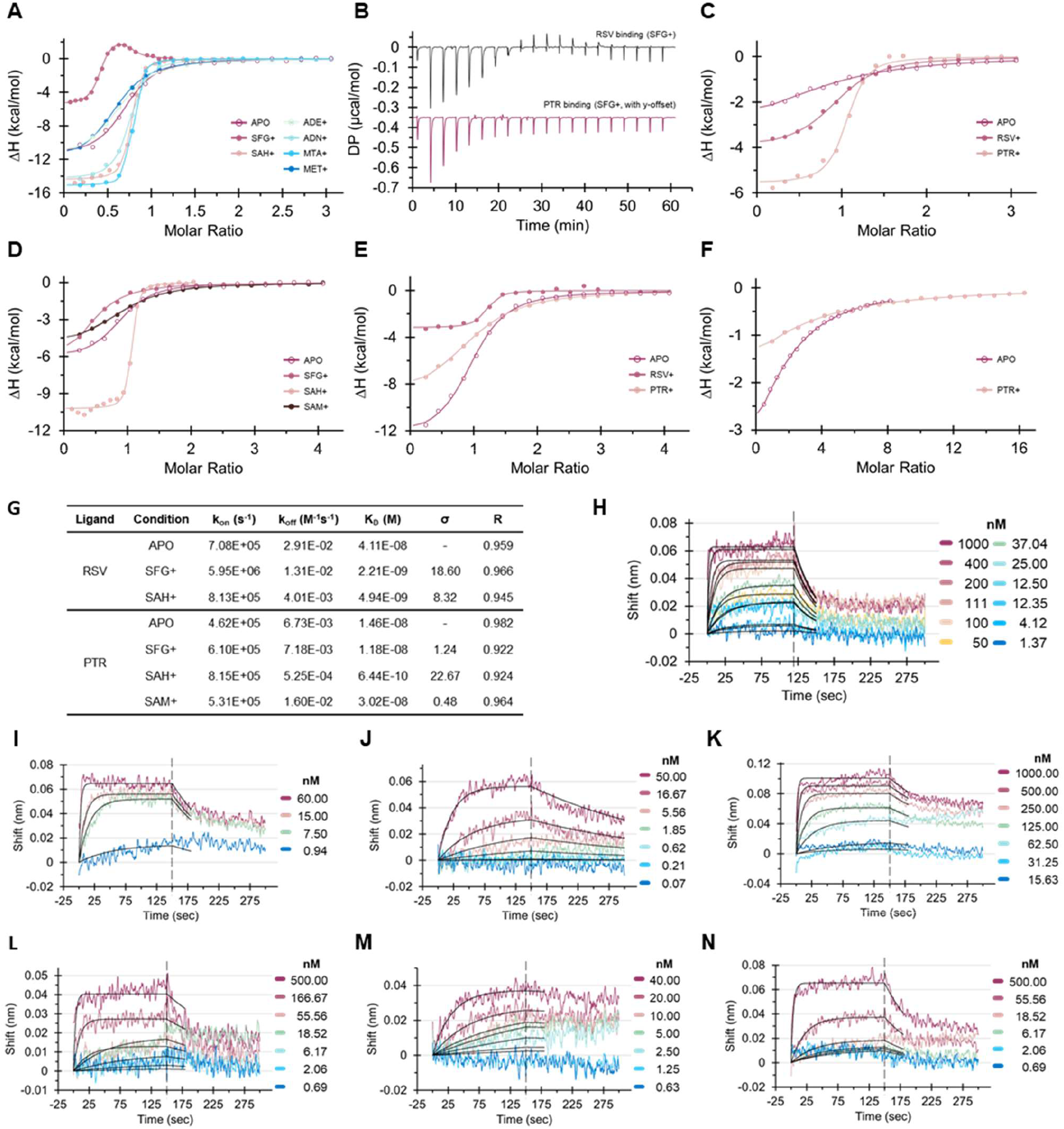
Differential cooperative bindings of SbSOMT towards cofactors and substrates. ITC plots of integrated heat peaks for the binding of towards RSV (**A**), SAH (**C**), PTR (**D**), SFG (**E**), and SAM (**F**) under apo (APO) condition or with 1 mM or saturating-condition of cognate-ligand (‘+’ marked) conditions. (**B**) Biphasic thermogram of the binding of RSV under SFG-saturating (SFG+) condition, in comparison to the monophasic thermogram of the binding of PTR under SFG+ condition. (**G**) Binding kinetics and cooperativity of SbSOMT measured by BLI. BLI sensograms of RSV binding under APO condition (**H**), SFG+ condition (**I**), and SAH+ condition (**J**). BLI sensograms of PTR binding under APO condition (**K**), SFG+ condition (**L**), SAH+ condition (**M**), and SAM+ condition (**N**).

The affinity profile towards SAH also exhibits a similar trend except both RSV-saturating and PTR-saturating conditions exhibit monophasic bindings (Fig. 1C). In particular, SAH affinity (675 nM) and positive cooperativity (σ of 37.78) are also stronger in PTR-saturating condition as compared to RSV-saturating condition (K_D_ of 3.96 μM, σ of 6.44). This observation suggests that SbSOMT is in favor of pairing cofactor and substrate that can fill the cavity at the active site with mass, such as the methyl group in PTR and SAM, and the amine group of SFG. The active site refers to the supposed cavity to accommodate the transferrable methyl group of SAM. Complementary to the loss of methyl group in SAH, the di-methylated PTR can contribute its methyl group to occupy the active site. This is further supported by the strong positive cooperativity for PTR binding in SAH-saturating condition (Fig. 1D) and SFG binding in RSV-saturating condition (Fig. 1E), which show σ of 35.26 and 8.88, respectively.

### Negative cooperativity in the unproductive pairing of SAM and pterostilbene

Attempt to crowd the active site with two methyl groups, however, negatively affects the cofactor-substrate cooperativity. The pairing of PTR to either SAM or SFG switches the cooperativity from positive to negative (Fig. 1, D to F). The negative cooperativity is relatively weak for the case SAM binding in PTR-saturating condition (Fig. 1F), where the affinity was weakened by σ of 0.81 to 139 μM; others show negative cooperativity ranging from 0.40 to 0.45. In particular, the stoichiometry was once again halved (N of 0.45) during the binding of PTR in SFG-saturating condition; attesting to the dimer asymmetry characteristic of SFG-bound SbSOMT. However, the titration was exhibiting a monophasic thermogram (Fig. 1B); implying only half of the substrate binding sites are available for PTR binding. In other words, the unoccupied half are available for the binding of reactive substrates (RSV or PNS). This dimer asymmetry derived from SFG-saturating conditions is however not inherently observed in SAM-saturating condition (N of 0.91). Nevertheless, we reason that the observed dimer asymmetry is in association with the conformational shift of SbSOMT rather than the sole act of SFG. Hence, the observed dimer asymmetry is likely an imbued nature of SbSOMT that was amplified in the SFG-binding scenario and might play role in the catalytic cycle.

In summary, the comprehensive binding affinity profile has unveiled that SbSOMT exhibits bilateral and selective positive cooperativity between cofactor and substrate. At the second binding event, the interplay at active site cavity plays a pivotal role in governing cooperativity switch; translating to meritorious intrinsic design for enzyme competency.

### Adenosine and 5-methylthioadenosine triggers cofactor-dependent positive cooperativity

The cooperativity effect of cofactor fragments was also examined to identify the capability of cofactor’s functional groups in triggering positive cooperativity. The affinity of RSV was examined via ITC in conditions consisting 1 mM of cofactor fragment (adenine (ADE), adenosine (ADN), 5-methylthioadenosine (MTA) or methionine (MET)). Intuitively, these conditions skew the SbSOMT population to fragment-bound state but not necessarily be saturating. Nevertheless, the results show that ADN (as the smallest functional group) and MTA are capable of triggering positive cooperativity (Table 1 and Fig. 1A). The findings indicate that the complete binding of cofactor backbone is unnecessary to trigger the positive cooperativity while highlighting the role of adenosine functional group for the following structural studies. At the same time, it ponders the physiological relevance of these cofactor fragments, which are essentially the ubiquitous metabolites, in manipulating the in vivo dynamics and activities of SbSOMT and methyltransferases in general.

### Insights of the conformational entropy correlated to SbSOMT cofactor-substrate cooperativity

The global thermodynamics recorded by ITC can provide insights to the cooperativity-associated conformational entropy (ΔS_conf_). Although the global thermodynamics is influenced by multiple factors (e.g., protein-ligand interaction, dilution of titrant, solvation), the apparent entropy (-TΔS) is often in good agreement to conformational entropy (*17, 18*), qualitatively to the very least. In particular, we find it worth to discuss the apparent entropy for the titrations of the RSV-SFG pair (Table 1) which has shown considerable thermodynamics flux by comparison.

The titration of RSV into apo SbSOMT, the first binding event, showed unfavorable entropy (-TΔS of 3.98 kcal/mol). Conversely, the asymmetrical endothermic and exothermic bindings at the second binding event were reactions with favorable entropies,-TΔS of-10.90 kcal/mol and-4.92 kcal/mol, respectively. This is not the case for SAH-bound SbSOMT which showed a slightly more unfavorable entropy (-TΔS of 5.26 kcal/mol) than the first binding event. Meanwhile, the apparent entropy of SFG binding at second binding events was reduced disproportionally from the first binding event (-TΔS of 5.43 kcal/mol). The titrations showed that the SFG binding towards RSV-bound SbSOMT was favorable (-TΔS of-5.41 kcal/mol) while the SFG binding towards PTR-bound SbSOMT remained unfavorable (-TΔS of 2.74 kcal/mol). Taken together, the discussed apparent entropies suggest that the SFG-RSV pair (and likely the SAM-RSV pair) prepays the entropic penalty for each other, leading to a favorable conformational entropy during the second binding event scenario.

### SFG and SAH trigger differential binding kinetics in attaining positive cooperativity

Biolayer interferometry (BLI) is an orthogonal method to unveil ligand binding kinetics through the measurement of association rate constants (k_on_) and dissociation rate constants (k_off_). The BLI analysis was successfully performed to resolve the binding kinetics of substrate analogues and the cofactor-dependent cooperativity. While the binding affinities measured by BLI were shifted proportionally to nanomolar range, the cooperativity characteristics were faithfully recapitulated (Fig. 1, G to N).

Strikingly, SFG and SAH demonstrated completely opposite effect on RSV binding kinetics to attain the positive cooperativity. In SFG-saturating condition, the k_on_ was accelerated greatly by 8.40 fold to 5.95 × 10^6^ M^-1^s^-1^, and k_off_ was decelerated by only 2.22 fold to 1.31 × 10^-2^ s^-1^. In SAH-saturating condition, the k_off_ was instead decelerated by 7.26 fold to 4.01 × 10^-3^ s^-1^ and k_on_ was accelerated by a mere 1.15 fold to 8.13 × 10^5^ M^-1^s^-1^. The SAH-driven binding kinetics show consistent trend in the case of PTR where the k_off_ was decelerated greatly by 12.82 fold in comparison to the 1.76-fold acceleration of k_on_; resulting in a positive cooperativity of 22.67. A negative cooperativity (σ of 0.48) comparable to the value determined by ITC is observed in SAM-saturating condition, where the binding towards PTR showed 2.38-fold acceleration of k_off_ in combination with a k_on_ comparable to apo condition. Meanwhile, SFG saturation drove slight positive cooperativity towards PTR (σ of 1.24) with proportional changes of k_on_ and k_off_.

In summary, the binding kinetics analysis of BLI has unveiled unprecedented discrepancy in the binding kinetics driven by SAM, SAH and SFG. The results imply that SAM binding can promote substrate association and product dissociation while SAH binding can prolong the residence time of both substrate and product. Yet, the enhancing effect of SFG on substrate k_on_ might be unfavorable in terms of the pharmacological effects. Collectively, the binding kinetic analyses have uncovered novel details of the cofactor-substrate cooperativity and interesting intrinsic properties valuable for methyltransferases’ enzyme and drug studies.

### Crystal structures of SbSOMT at different states unveil dynamical SAM-binding domain and binding basis of SAH

Previously, our group had solved the crystal structures of SbSOMT in complex with substrate analogues to decipher the structural basis of SbSOMT substrate recognition (*16*). Here, the crystal structures of apo SbSOMT and SAH-bound SbSOMT were solved and compared altogether with RSV-bound structure (PDB: 7VB8) to elucidate the dynamic transitions of SbSOMT across different states and address the underlying mechanism of positive cooperativity.

The apo SbSOMT was solved at 2.04 Å with a large asymmetric unit encompassing three dimeric SbSOMT (fig. S2A). All six chains appear in open conformation where the SAM-binding domain is opened and the substrate binding pocket is accessible (Figs. 2, A and B, and S2B). In particular, the crystal packing lacks crystal contacts around the SAM-binding domain of Chain E and Chain F (Fig. 2C), resulting in the wider opening and unstable conformation of SAM-binding domains (fig. S2B). By comparison, these domains exhibit above average B factor (Fig. 2D) and relatively poor electron density map (fig. S2C). It is worth noting that the crystal structure has demonstrated potential higher-order structures of apo SbSOMT. In the crystal structure, a symmetrical dimer-of-dimer tetramer is observed where the dimers inversely clamp the opponent N-terminal loop (69-76) via SAM binding domain opening (fig. S2D). Additionally, a partial inter-dimer β-sheet is formed between 6^th^ β-strand of SAM binding domain Rossmann fold core (361-367) and the transient β-hairpin (96-104) of the N-terminal core (fig. S2E).

**Fig. 2.**
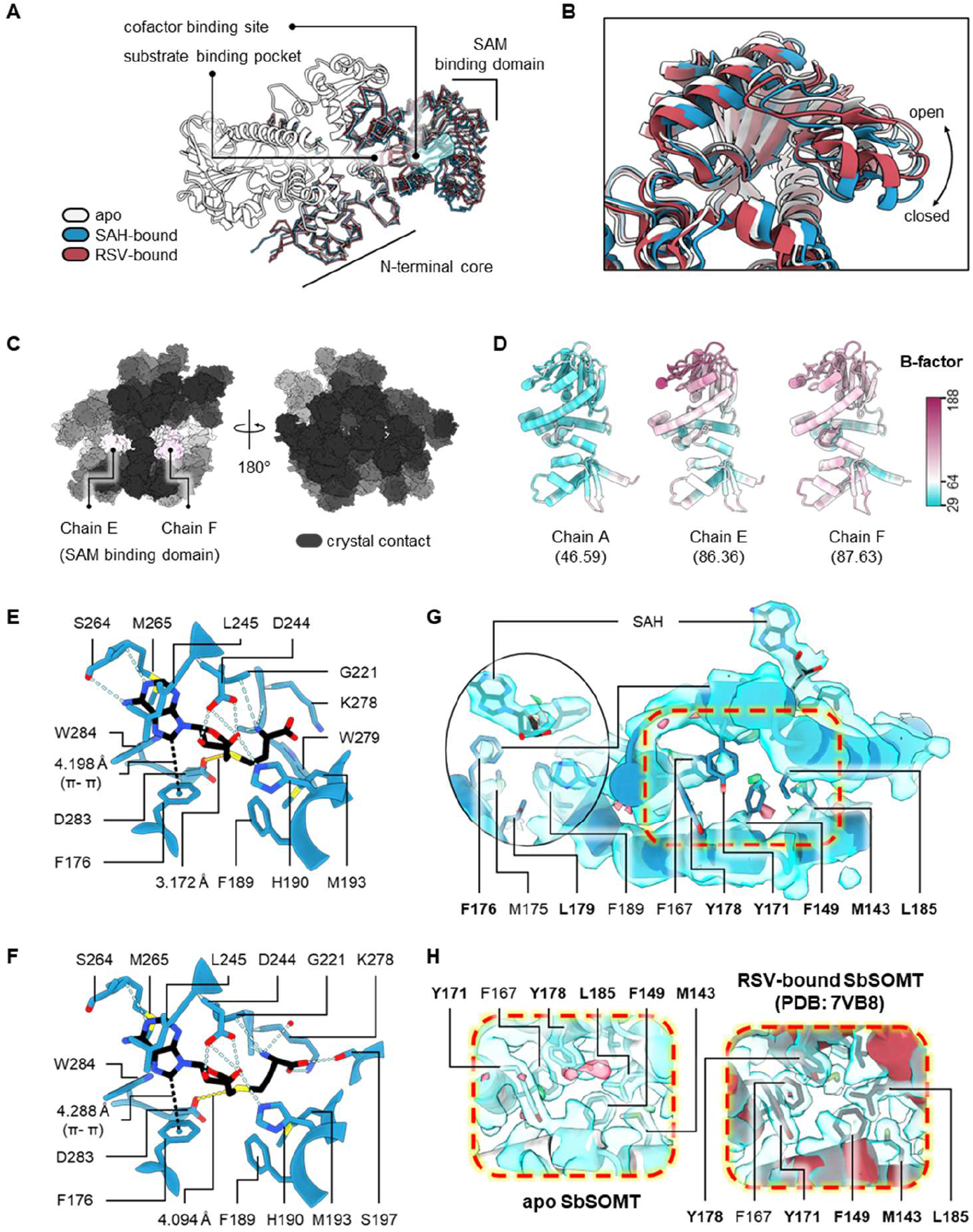
Structural insights of apo SbSOMT and SAH-bound SbSOMT. (**A**) Overall architecture of SbSOMT and superimposition of SbSOMT in apo (white), SAH-bound state (blue), and RSV-bound state (red, PDB: 7VB8) (*16*). (**B**) The SAM binding domain exhibits different conformation in apo (open), RSV-bound (partially-closed) and SAH (closed) states. (**C**) SAM binding domain of Chain E and Chain F lacks crystal contacts (≤ 3 Å, black) in apo SbSOMT. (**D**) In apo SbSOMT, the B-factor of Chain E (86.36) and Chain F (8.63) are above average (64.0) and localized majorly at SAM binding domain. SAH binding basis of Chain A (**E**) and Chain B (**F**) of SAH-bound SbSOMT features F176-adenine π-π stacking (black dotted line), varied D283-sulfur-center distance (yellow dotted line), and varied network of hydrogen bonds (blue dotted line). (**G**) Electron density maps of SAH-bound SbSOMT reveals poor density and unstable side chain of hydrophobic residues (labels in bold) at SAH proximity (squared by red dotted line). Conversely, these residues exhibit clear density and stable conformation in apo SbSOMT (**H**, left) and RSV-bound SbSOMT (**H**, right). The electron density maps (**G, H**) are 2Fo-Fc map contoured at 1.0σ (blue surface) and Fo-Fc maps contoured at 3.0σ/-3.0σ (green/red surface).

On the other hand, the SAH-bound SbSOMT binary complex was solved at 2.90 Å, exhibiting a closed conformation (Figs. 2B and S3, A and B). In association with the binding of SAH, SbSOMT exhibits tighter closure of SAM binding domain than the previously reported RSV-bound structure (Fig. 2B). SAH is bound to both protomers and blocking the substrate pocket, albeit with subtle variations (Figs. 2, E and F, and S3C). In both chains, the adenine moiety is sandwiched by L245 and M265; the ribose moiety forms the conserved bidentate bond with D244 and an additional hydrogen bond with H190. In Chain A, the distance between SAH sulfur center and D283 is fitted at 3.172 Å, ∼0.9 Å closer than the case in Chain B. The SAH of Chain B is instead stretched by the hydrogen bonds at the amide end, between the carboxylic group and rotamers of S197 and K278. Meanwhile, the hydrophobic patch (M143, F149, Y171, F176, Y178, L179, L185) at SAH proximity is unstable, where the residues compartmentalizing the hydrophobic substrate pocket exhibits poor density resolution (Fig. 2G). The destabilization of the hydrophobic network is likely associated with the π-stacking between F176 and adenine moiety of SAH (Fig. 2, E and F). The hydrophobic patch of apo SbSOMT and RSV-bound SbSOMT with free F176 exhibits well-resolved density and stable conformation (Fig. 2H). On the other hand, the interaction between F176 and adenine is conserved and critical for enzyme activity of plant *O*-methyltransferase, COMT-S (*19*). Structurally, few homologs have showcased shorter length of the F176-adenine π-π interaction (*20, 21*) (fig. S4, A and B); while few others showcase a flexible and unmodelled hydrophobic patch (*22, 23*) (fig. S4, C and D). Taken together, the unstable hydrophobic patch of SAH-bound SbSOMT might serve as a strategic opening for substrate binding, in compensation to the tightening of SAM binding domain and the blockage of cofactor.

### Integrative insights reveal W279 π-stacking network governs cofactor-substrate cooperativity of SbSOMT

Integrating the structural insights across different states of SbSOMT further reveal the transitional dynamics of SbSOMT from apo state to the two ligand-bound states (Fig. 3, A and B). In good agreement with the results of binding kinetics, SbSOMT exhibits different conformational landscapes between first and second ligand binding events.

**Fig. 3.**
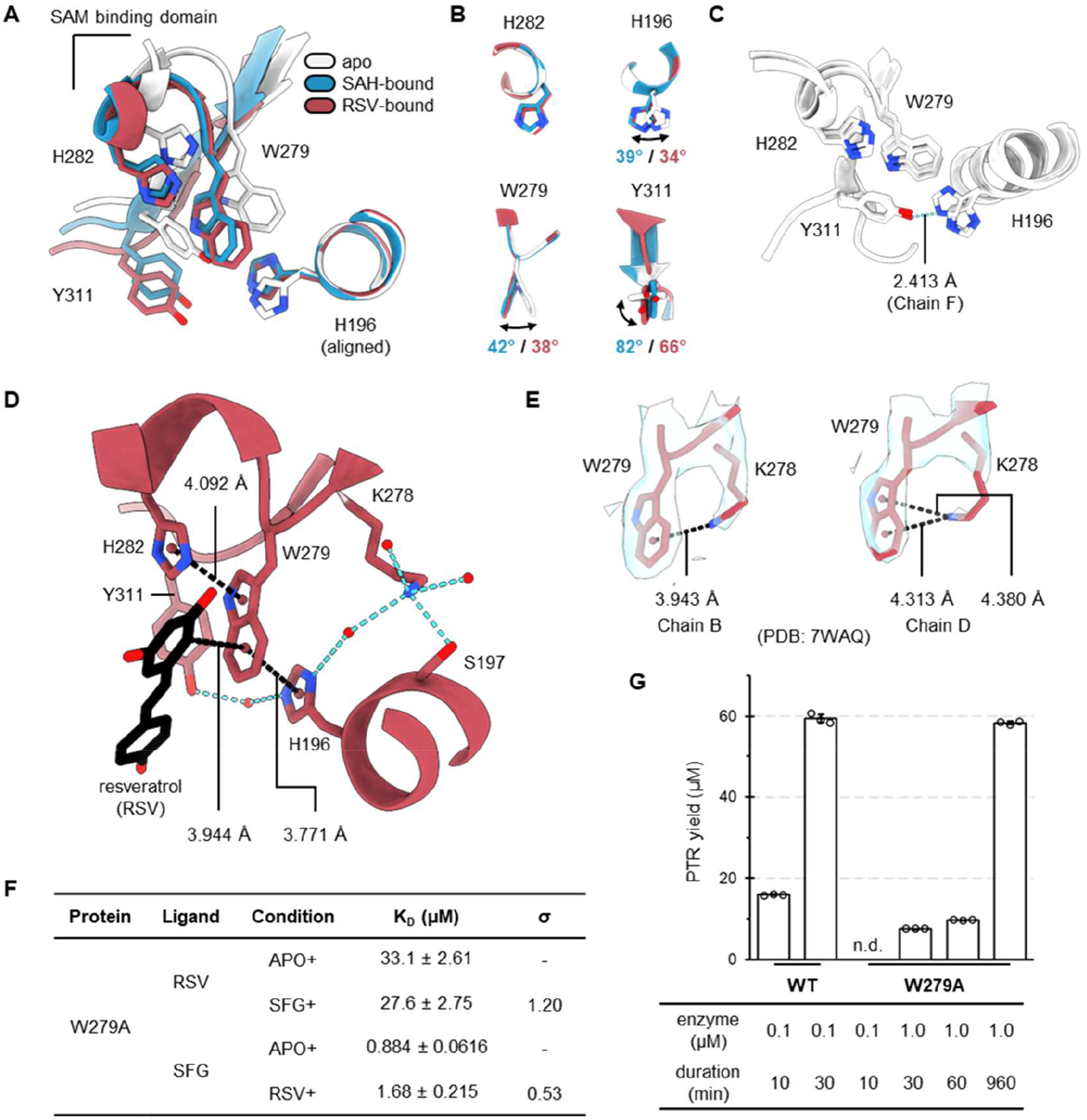
Structural basis of dynamical W279 π-stacking network governing cofactor-substrate cooperativity of SbSOMT. (**A**) Superimposition at H196 reveals the domain and rotamers rearrangement at W279 proximity. W279 is shielded by H196 and Y311 in apo state (white), but exposed in SAH-bound (blue) and RSV-bound (red, PDB: 7VB8) states. (**B**) Comparison of rotamer conformation of H196, H282, W279 and Y311 from different states. (**C**) H196-Y311 hydrogen bond (blue dotted line) is only observed in Chain F of apo SbSOMT. (**D**) The π-stacking network represented by RSV-bound SbSOMT reveals that W279 indole ring can form three π-stacking interactions (black dotted line), either with H196, H282 or RSV. Blue dotted lines depict the hydrogen bonds. (**E**) Alternative RSV-bound SbSOMT crystal structure (PDB: 7WAQ) demonstrates K278 flexible conformations and capability to form the fourth π-stacking interactions towards W279. The 2Fo-Fc map contoured at 1.0σ (blue surface). (**F**) ITC binding affinity profile of W279A mutant demonstrates the loss of cooperativity in the mutant. (**G**) Comparison of PNS-methylating enzyme activity of wildtype SbSOMT (WT) and W279A mutant demonstrate significant loss of activity in the mutant. PTR yield of the 10 min reaction of W279A mutant was not detected (n. d.) in the HPLC analysis.

**Fig. 4.**
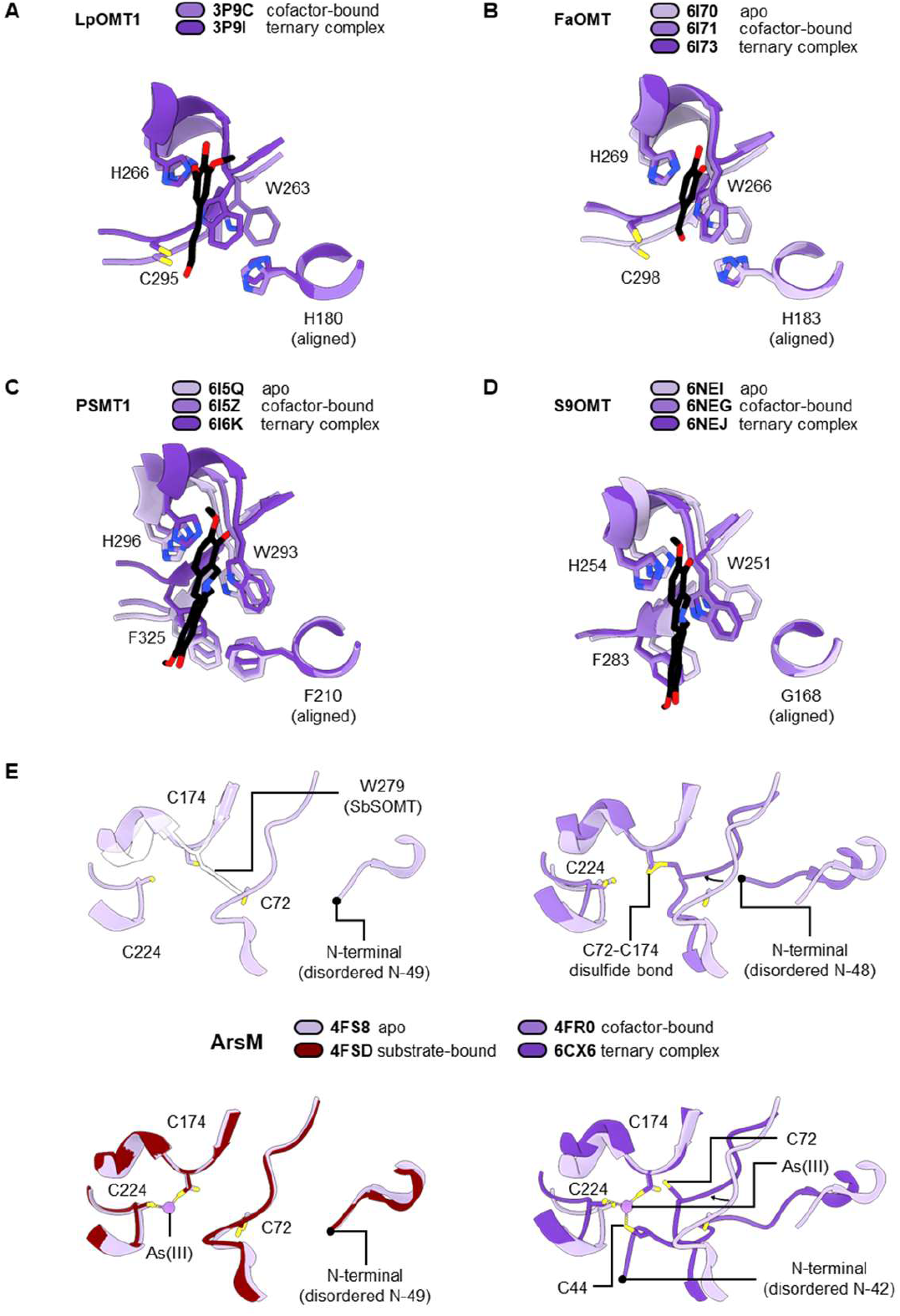
Analysis of conformational dynamics in Class I SAM-dependent methyltransferases and potential cofactor-substrate cooperativity. (**A** to **D**) Retrieved structures of W279-conserved plant *O*-methyltransferase homologs that are deposited in multiple states. Each legend denotes the PDB IDs and deposited states of the methyltransferase. Residues with shown side chain and label (in sequential order) are positioned equivalent to H196, W279, H282 and Y311 of SbSOMT, respectively. Ligand (black backbone) refers to substrate analogs bound to the ternary complex. The caffeic acid *O*-methyltransferase pairs (**A**, LpOMT1 (*27*) and **B**, FaOMT (*20*)) and scoulerine 9-*O*-methyltransferase pairs (**C**, PSMT1 (*21*) and **D**, S9OMT (*28*)) show no consensus in the shifting of tryptophan and state transition. Only FaOMT and S9OMT demonstrate pre-alignment of tryptophan in cofactor-bound state for ternary complex formation. (**E**) Structural analysis of arsenite methyltransferase (ArsM) across different states (*35-37*). The legend denotes the PDB IDs and deposited states of ArsM. ArsM conserves four cysteine (C44, C72, C174 and C224) at active site proximity where C174 is equivalent to W279 of SbSOMT. C174 can form C72-C174 disulfide bond, and coordinate arsenite, As(III), binding together with C44 and/or C224. ArsM exhibits different degrees of conformational shift (curved arrow) and N-terminal disorder in different states.

At apo state, the SAM binding domain exhibits an open conformation where W279 is shielded by H196 and Y311 from exposing to the binding pocket (Fig. 3C). In which, Chain F further show a hydrogen bond between H196 and Y311. In both SAH-bound and RSV-bound structures, the SAM binding domain exhibits a closed conformation and W279 is exposed to form a π-stacking network (Fig. 3, A and D). In other words, the transition from apo state to SAH-bound or RSV-bound states triggers the closure of SAM binding domain and local rearrangement at W279 proximity. The exposed W279 indole ring serves as a central plane feasible for four π-stacking interactions. As exemplified with RSV-bound SbSOMT (Fig. 3D), the pyrrole ring stacks with catalytic H282 and the benzene ring stacks with the C2 of substrate at the exposed facade; essentially facilitating the interaction between NE2 of H282 and 3-OH of RSV. At the rear of the plane, the benzene ring stacks with H196. In another RSV-bound structure (PDB: 7WAQ), a transient π-stacking is formed by the flexible K278 towards W279 rear plane (Fig. 3E).

The π-stacking network is associated directly and indirectly to the closure of SAM binding domain. The H196-W279 π-stacking directly bridges the H196-attached N-terminal core with the W279-attached SAM-binding domain. Meanwhile, the bound substrate indirectly contributes to the domain closure as it sits in the binding pocket which is majorly constituted by N-terminal core, while establishing interactions with SAM-binding-domain-anchored H282 and W279. In short, the evident W279 π-stacking network and the transient H196-Y311 interactions represent the significant energy barrier that differentiate the conformational landscapes at first and second ligand binding events.

The integrative insights imply that either the binding of substrate or cofactor can overcome the energy barrier and reshaping the conformational landscapes to favor the consecutive binding of the cognate ligand and subsequent formation of catalytic ternary complex. Taken together, we propose the structural mechanisms that drive positive cooperativity in SbSOMT based on W279 π-stacking network. For the substrate-dependent cofactor cooperativity, the binding of substrate leads to the formation of W279 π-stacking network. The re-oriented W279 does not involve directly in SAH binding but in turn, contributes to the closure of SAM binding domain and bringing close the cofactor-interacting residues at N-terminal for π stacking (F176), hydrogen bonds (H190 and S197) and hydrophobic interactions (F176, F189 and M193). Notably, our previous study has observed that the RSV-bound SbSOMT was prone to co-crystallize with NAD^+^ (PDB: 7VB8, (*16*)) where the adenyl-moiety is stably stacked between Leu245 and Met265. In other words, the binding of substrate triggers the induced-fit of cofactor binding site to better accommodate the cofactor.

For the cofactor-dependent substrate cooperativity, the binding of cofactor forms adenine-F176 π-stacking and leads to the pre-alignment of W279 π-stacking network and destabilization of the adjacent hydrophobic patch (M143, F149, Y171, F176, Y178, L179, L185). The destabilized patch increases the exposure of the binding pocket and the previously-buried W279 benzene façade is essential for substrate binding. These unstable hydrophobic side chains are essentially in favor towards the binding of substrate with hydrophobic stilbene backbone over the continuous exposure towards the hydrophilic bulk solvents. As a result, the cofactor-bound SbSOMT show greater favorability towards substrate binding as compared to apo SbSOMT. Meanwhile, we speculate that the instability of hydrophobic patch is linked to the species of bound cofactor, leading to the differential binding kinetics observed in BLI analyses. For instance, the protrusion and electrostatic of SFG’s amine group intensify the instability of the hydrophobic patch and enhance the k_on_ of substrate.

To further validate the role of W279 π-stacking network, W279 was substituted with alanine and the mutant’s binding and enzymatic activities were examined. The binding affinity profile of W279A mutant evidently indicate that the loss of positive cooperativity and significant impairment on the RSV affinities (Fig. 3F). In SFG-saturating condition, the RSV binding depicted a monophasic thermogram and the K_D_ (27.6 μM) was comparable to apo condition (33.1 μM). In contrast, W279A affinity towards SFG was stronger in apo condition (0.88 μM) and comparable to RSV-bound SbSOMT. The impaired RSV affinity reflects the C2-W279 π-stacking interaction contribute critically in substrate binding. Expectedly, the enzymatic activity of W279A mutant was also significantly impaired (Fig. 3G). W279A reaction consumed 10-fold more enzyme concentration and 32-fold longer duration to achieve a comparable yield as compared to wildtype. Both binding and enzymatic activities of W279A mutant confirm that W279 π-stacking network governs SbSOMT positive cooperativity.

## Discussion

In this study, we have discovered and characterized the bilateral and selective positive cofactor-substrate cooperativity of SbSOMT, unknown to literature. Our finding elaborates the unprecedented and delicate interplay between methyltransferase with cofactor, substrate, product and inhibitors. The cofactor-substrate cooperativity is positive and bilateral in general. The positive cooperativity permits activation by few partial functional groups of the cofactor but is intolerant to the steric hindrance at the active site cavity. The steric hindrance leads to the cooperativity switch in the binding kinetics of SAM-PTR and SFG-PTR pairs but not SAH-PTR pair. This correlation suggests SAM exhibiting SFG-like outcomes such as enhanced positive cooperativity with favorable conformational entropy and competent binding kinetics towards RSV binding. Particularly, SFG-bound SbSOMT demonstrated positive cooperativity via the disproportional acceleration of k_on_ while SAH led to the disproportional deceleration of k_off_ to achieve the comparable cooperativity. Integrative structural insights from SbSOMT at different states further pinpoint the role of W279 π-stacking network in governing SbSOMT cooperativity. The binding of the first ligand leads to the exposure of W279 and formation of the W279 π-stacking network along with the closure of the SAM binding domain. The W279 π-stacking network represents a significant energy barrier between the conformational landscapes with different ligand binding kinetics. Altogether, our study offers an in-depth understanding to the intrinsic structural dynamics governing SbSOMT cofactor-substrate cooperativity.

The observed features of SbSOMT cofactor-substrate cooperativity have reflected the enzyme’s intrinsic designs for catalytic competency. The merits of cooperativity selectivity are not limited for the discrepancy between di-hydroxy substrate and di-methylated product, but also creditable for recognizing 5’-hydroxy from 3’-methoxy in PNS intermediate. This overcomes the challenge of stilbene backbone symmetry that might compromise catalytic efficiency via unproductive binding orientation. Meanwhile, the shown accelerated RSV association rate of SFG-bound SbSOMT imply that SAM binding accelerates consecutive binding of reactive substrate, proactively promoting the formation of reactive ternary complex and shortening the window of binary complex state. At the same time, SAM was shown to accelerate PTR dissociation by near 2.5-fold, freeing the substrate pocket for reactive substrates. Conversely, the slowed substrate dissociation caused by SAH can be translated as a strategy to prolong the residence time of substrate in the binding pocket for the shift encounter with reactive SAM once the cofactor is exchanged. Nevertheless, stronger deceleration effect of SAH towards PTR dissociation can be a limiting step in SbSOMT catalytic cycle.

From drug design perspective, the contrasting effects of SFG and SAH towards substrate binding kinetics highlight the strong role of pharmacophore interacting with the active site. Despite SFG being the common cofactor-analogous inhibitor, SFG effects on accelerating RSV association and promoting substrate-bound state are the potential shortcomings from a pharmacological perspective. While SAH-PTR binding kinetics showcases an alternative in filling the active site cavity with meritorious prolonged residence time. In short, the active-site-interacting pharmacophore can be harnessed for binding kinetics manipulation in the drug designs for the common, cooperative or bi-substrate inhibitors.

The mechanism of cofactor-substrate cooperativity of SbSOMT represents the intrinsic and dynamical regulatory logic of enzymes that remains the enigmatic and understudied learning ground. The revolutionary AI advancements in structural biology and enzyme design have progressed but limited implications in the realm of structural dynamics. The leading enzyme design studies have accredited the importance in accurate design of active site constellations and transition states as major contributor in besting the design or as the major bottleneck in overcoming the catalysis turnover number (*24, 25*). Recent work on the emulation of protein conformational ensembles also marks the limitation owning to the lacking of high quality training data (*26*). Novel and dynamical structure-function relationships such as cooperativity, are still largely unannotated and overlooked; demanding experimental discovery and validation.

Exploring the protein conformational landscape at multiple states is a classic but robust approach to decipher the underlying dynamics. Though W279 was previously reported to reorient upon substrate binding (*27*), no literature has linked W279 to the π-stacking network and cofactor-substrate cooperativity. Most plant *O*-methyltransferases (InterPro: IPR016461) are deposited in single or invariant states with limited insights for structural dynamics investigation. In a non-exhaustive PDB search, we could only identify four sets of W279-conserved homologs with multi-states structures (Fig. 5, A to D); feasible for analyzing the potential cofactor-dependent cooperativity. They include two caffeic *O*-methyltransferases, LpOMT1 (*27*) and FaOMT (*20*), and two scoulerine 9-*O*-methyltransferases, PSMT1 (*21*) and S9OMT (*28*). Both caffeic *O*-methyltransferases exhibit the equivalent tryptophan π-stacking network for substrate interaction, yet only FaOMT shows a pre-aligned π-stacking network upon SAH binding. Similarly, PSMT1 and S9OMT do not share consensus trend on tryptophan rearrangement; only S9OMT reorients the tryptophan upon SAH binding.

Towards distal domain homologs (Class I methyltransferases), several human methyltransferases (PRMT1, PRMT5, hPNMT, RNMT, and etc.) are reportedly exhibiting positive cooperativity or cofactor-dependent sequential binding of substrate (*29-32*). Structural alignment with SbSOMT shows that these homologs do not conserve W279 (fig. S5) and lack multi-states structures to address the underlying cooperativity mechanism (*2, 32-34*). Intriguingly, we came across a set of arsenite methyltransferase (ArsM) structures that show a potential cooperativity network manifested by cysteine disulfide bond (Fig. 5E). As reported in the studies led by Rosen’s group (*35-37*), ArsM comprises four cysteine residues (C174 as W279 equivalent, C44, C72, C224). C174 can form cysteine disulfide bond towards C72 and introduce conformational shift upon cofactor binding; as well as coordinate the As(III) substrate together with C44 and/or C224. In sum, the lack of consensus in terms of rotamer dynamics and residues’ identity imply that the W279 π-stacking network in SbSOMT is an uncommon positive epistasis that can be harnessed to drive evolution of other homologs or designer enzymes.

Our study has demonstrated a novel mechanism of cofactor-substrate cooperativity in SbSOMT at structural basis. The W279 π-stacking network is but a limited representation to the structurally diverse SAM-dependent methyltransferases. Expanding the study to more representations will not only benefit the fundamental understanding and applications towards individual methyltransferases but pave way to the understanding of methyltransferases activity at higher order systems.

## Materials and methods

### Bacterial strain and culture conditions

*Escherichia coli* strains were cultured in 2xYT medium (16 g/L tryptone, 10 g/L yeast extract, and 5 g/L NaCl) comprising appropriate amount of antibiotics in a 37 °C orbital shaker, in general. *E. coli* DH5α was used for molecular cloning and sub-cloning purpose while *E. coli* BL21-CodonPlus (DE3)-RIL was used for the purpose of recombinant protein overexpression.

### Molecular cloning, recombinant protein overexpression, and purification

Plasmid construct, pH_6_SbSOMT, was constructed in previous study (*16*), comprising pET-N-His-TEV vector backbone and *SbSOMT*(2-377). Mutagenesis of SbSOMT, namely W279A mutant, was constructed via ClonExpress MultiS One Step Cloning Kit (Vazyme) using single PCR amplicon from designated primers (<u>W279A</u> Forward Primer:5’-GCAGTTCTGCTCAA<u>GGC</u>GATTCT TCATGATTGGGACG-3’; <u>W279A</u> Reverse Primer:5’-CTTGAGCAGAACTGCATTTC-3’).

The overexpression of recombinant wildtype SbSOMT or W279A mutant was conducted with slight modifications to previously reported protocol (*16*). Briefly, the overexpressing strain was freshly prepared from chemically-induced competent cells. Transformed cell was subcultured into 2xYT medium supplemented with kanamycin and chloramphenicol, and recovered at 37 °C via overnight shaking. Fresh 2xYT medium was inoculated with the overnight culture and shaken at 37 °C until the OD_600nm_ reached 0.6-0.8. Protein expression was then induced via 0.1 mM Isopropyl-β-thiogalactopyranoside (IPTG) at 16 °C, overnight.

Harvested cells was resuspended in Lysis Buffer (50 mM Tris, 150 mM NaCl, 50 mM glutamate, 50 mM arginine, 10% glycerol, 10 mM TCEP, pH 7.9) and lysed by high-pressure cell disruption system. The cell lysate was then centrifuged and clarified with 0.40 μm filter. Clarified lysate was loading into HisTrap HP column (Cytvia), followed by washing with 10 column volumes of Washing Buffer (50 mM Tris, 150 mM NaCl, 25 mM imidazole, 5 mM TCEP, pH 7.9). The bound protein was then eluted by imidazole gradient buffered by the buffer (50 mM Tris, 150 mM NaCl, 5 mM TCEP, pH 7.9). The eluted fractions containing desired protein were pooled and mixed with TEV protease. The mixture was concentrated and exchanged to Low Salt Buffer (20 mM HEPES, 50 mM NaCl, 1 mM TCEP, pH 7.9) by ultrafiltration, followed by overnight incubation at 4°C. The incubated sample was clarified and flowed through a HisTrap column prior to loading into HiTrap Q HP column (Cytvia) for anion exchange chromatography. The loaded column was washed with Low Salt Buffer until the baseline had been stabilized. Salt gradient up to 1.0 M was employed to elute the bound protein. Desired fractions were pooled and proceeded for ultrafiltration. The protein stock was equilibrated to Storage Buffer (10 mM HEPES, 100 mM NaCl, pH 7.9) and flash frozen by liquid nitrogen for long-term storage at-80°C.

### Isothermal Titration Calorimetry (ITC)

ITC binding experiments were conducted using MicroCal PEAQ-ITC instrument (Malvern). The running buffer of ITC assays comprised 100 mM HEPES (pH 7.9) and 0-5.0 v/v% PEG 400 (as specified in table S2). The concentrations of protein (in cell), ligand (in syringe) and saturating-ligand (constant, in both cell and syringe) that were used in each run are listed in table S2. The stock proteins were stored at least 10-fold concentrated as compared to the final concentration (in cell) in Storage Buffer. The stocks of substrate analogs were dissolved in 50 v/v% PEG 400 aqueous solution. Except SAH (50 mM stock in 50 mM HCl), SFG and SAM stocks were dissolved in water.

All ITC runs were conducted at 25 °C, 750 rpm stirring speed, and with reference power of 10.0 μcal/s. Each run was initiated with 60 s delay, followed by a 0.5 μL first injection and consecutive 19 injections of 2.0 μL. The intervals between injections were all 180 s. The thermograms were analysed using MicroCal PEAQ-ITC Analysis Software (Malvern). Except for one exception, the results were calculated by integrating the injections from 2^nd^ to 20^th^ injection, using the “One Set of Sites” fitting model at default settings. For the ITC run of RSV binding in SFG-saturating condition, the results were calculated by integrating the 18 injections and excluding 1^st^ and 15^th^ injection, using the “Two Set of Sites” fitting model at default settings. The individual thermograms and integrated plots are available in figs. S6 and S7.

### Calculation of cooperativity (σ)

Cooperativity (σ) can be calculated based on the binding affinity of ligand at first binding event (apo condition) over the binding affinity of ligand at second binding event (cognate-ligand-saturating condition). The equation is listed as follow:

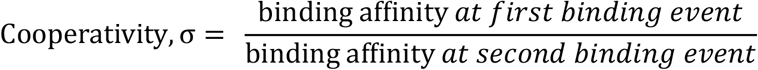

In which, σ > 1.0 implies positive cooperativity; σ = 0 implies no cooperativity; σ < 1.0 implies negative cooperativity.

### Biolayer Interferometry (BLI)

BLI binding experiments were conducted using Gator Prime instrument (Gator Bio) and SA XT probes (Gator Bio) at 30 °C. Briefly, SbSOMT was first biotinylated by EZ-Link NHS-LC-LC-biotin (Thermo Fisher Scientific) following manufacturer’s protocol, and then equilibrated in Loading Buffer (20 mM HEPES, 150 mM NaCl, pH 7.4). The probes were pre-wetted with Loading Buffer and the biotinylated SbSOMT was loaded onto the probes via hour-long incubation until the probes were fully saturated. The loaded probes were washed via incubation in a new Loading Buffer until a stable baseline was observed. The prepped probes were then proceeded for BLI assay.

Each BLI assay consists of three steps – baseline (60 s), association (120 or 150 s) and dissociation (180 s). In all steps, the assay buffer comprised 20 mM HEPES, 150 mM NaCl, 1% DMSO, at pH 7.4, as the basic composition. In SFG or SAH-saturating conditions, the basic composition of the assay buffer in all three steps was topped with 50 μM of SFG or SAH, respectively. Lastly, the buffer composition in association step comprised additional ligand of interest (RSV or PTR) with a concentration gradient (listed in figure legend) across different wells. The sensograms were screened and analyzed using GatorOne software (Gator Bio). Sensograms generated by GatorOne is available in fig. S8.

### Protein crystallization

Initial crystallization screenings of apo SbSOMT (10 mg/mL SbSOMT, 10 mM HEPES, 100 mM NaCl, pH 7.9) and SAH-bound SbSOMT (10 mg/mL SbSOMT, 2 mM SAH, 10 mM HEPES, 100 mM NaCl, pH 7.9) were first conducted using commercial kits (Hampton Research and QIAGEN). The screenings were conducted in sitting drop vapor diffusion fashion where 100 nL protein sample and 100 nL crystallization reagent were dispensed together by Mosquito liquid handler (TTP LabTech), and incubated at 18 °C. Crystallization of apo SbSOMT (20 mg/mL SbSOMT, 10 mM HEPES, 100 mM NaCl, pH 7.9) was optimized in crystallization reagent comprising 0.2 M sodium malonate (pH 7.0), 20% PEG 3350, and 10% glycerol. Equal volume (1.0 μL) of protein and crystallization reagent were mixed together and sealed with 100 μL reservoir. The crystals were obtained after >4 weeks incubation at 18 °C. Co-crystallization of SAH-bound SbSOMT (30 mg/mL SbSOMT, 20 mM HEPES, 100 mM NaCl, pH 7.9) was optimized in crystallization reagent comprising 0.04 M citric acid/0.06 M bis-tris propane (pH 6.4), 20% PEG 3350; and 5% glycerol. The drop was automatedly dispensed by the sequential addition of 500 nL protein, 250 nL ligand (50 mM SAH in 50 mM HCl), and 500 nL crystallization reagent. The crystals were obtained after ∼1 week incubation at 18 °C. The crystals were harvested and cryoprotected by 20% glycerol. The cryoprotected crystals were mounted on a loop, followed by cryo-cooling with liquid nitrogen. The vitrified crystals were stored in a puck under cryo-condition until data collection.

### X-ray crystallography

Single crystal cryo-crystallography was employed to obtain X-ray diffraction datasets of the crystals. The datasets were collected from beamline BL02U1 (*38*) of Shanghai Synchrotron Radiation Facility (SSRF), Shanghai, China. The raw data were processed by XDS (*39*) using the autoproc automated pipeline(*40*). CCP4 suite (*41*) was used in the following steps. The unmerged data was reduced by Aimless (*42*). Molecular replacement was conducted via PHASER (*43*) to solve both apo and SAH-bound SbSOMT via previously published structure (PDB: 7VB8, (*16*)). Refinement of the structures were performed automatedly via REFMAC5 (*44*) and PDB-REDO (*45, 46*), and manually via COOT interface (*47*). Compilation of structure factor and coordinate of final deposition, and generation of data collection statistics were facilitated by OneDep (*48*). The statistics and metrics of reported structures are compiled in table S3. Figures featuring protein structures were prepared using UCSF ChimeraX (*49*).

### In vitro enzymatic assay

Methylation activity of purified wildtype SbSOMT and W279A mutant were examined using varied final enzyme concentrations (0.1 and/or 1.0 μM) and varied incubation durations (10, 30, 60 and/or 960 min) with 0.5 mL final reaction mix comprising 0.1 mM HEPES (pH 7.9), 0.5 mM SAM, 0.5 mM PNS, and 10% methanol. Reactions were conducted at 30 °C for a determined duration, followed by quenching via ice and equal volume of acetonitrile.

The enzymatic activity was determined by quantifying the PTR product via high performance liquid chromatography (HPLC) on a HPLC system equipped with Diode Array Detector (Agilent) and C18 column (YMC-Pack ODS A, YMC). Briefly, the quenched mixture was first filtered through a 0.22 μm PVDF filter prior to loading. The analyte (10 μL) was loaded into a pre-equilibrated column with mobile phase of 70% water and 30% methanol. After loading, the mobile phase was gradually increased to 100% methanol over a course of 6 min at 0.5 mL/min flow rate, followed by 3 min of isocratic mobile phase (100% methanol) at the same flow rate. PTR was detected based on UV absorption at 334 nm. Triplicates were performed for each enzymatic reaction. The chromatograms (figs. S9 and S10) were processed via Agilent OpenLab software. Quantification of PTR was conducted using a calibration curve of PTR standards (fig. S11).

## Acknowledgements

We thank Shenzhen Synthetic Biology Infrastructure (InfraSynBio) and Core Facility of Shenzhen Institute of Synthetic Biology (iSynBio) for providing ITC and HPLC facilities. We thank Shanghai Synchrotron Radiation Facility (SSRF) and BL02U1 (SSRF) for providing X-ray crystallography beamtime and BL02U1 beamline scientists for technical support. This work is funded by Basic and Applied Basic Research Foundation of Guangdong Province, China (Grant No.: 2023B0303000003) and Research Grants Council of Hong Kong, China (Grant No.: GRF17111423). K.C.P. acknowledges the support of Hong Kong PhD Fellowship Scheme during his affiliation with HKU, and the support of Guangdong Province Overseas Postdoctoral Talent Support Program during his affiliation with Spallation Neutron Source Science Center and Institute of High Energy Physics.

## Notes

### Competing Interest Statement

The authors have declared no competing interest.

