## Supplementary Materials for "Structural basis of the π-stacking network governing cofactor-substrate cooperativity of SbSOMT"

**This PDF file includes:**

Figs. S1 to S11

Tables S1 to S3

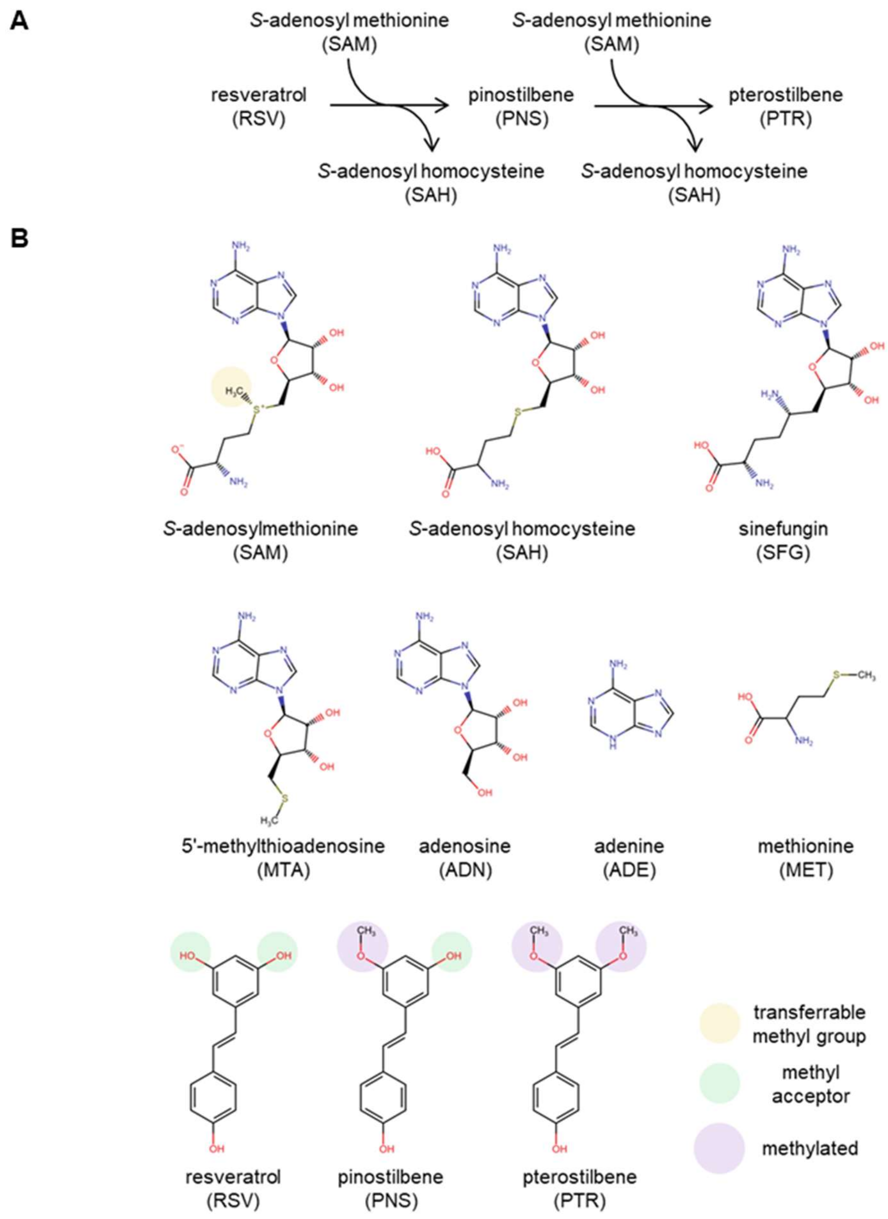


**Fig. S1. SbSOMT catalytic activity and analogs of cofactor and substrate.** (**A**) Schematic view of SbSOMT catalytic activity. (**B**) Chemical structures of cofactor and substrate analogs used in this study. The transferrable methyl group (yellow), methyl acceptor (green) and methylated site (purple) are circled. The chemical structures are generated using Marvin JS by Chemaxon (https://www.chemaxon.com) available on RCSB portal (https://www.rcsb.org/chemical-sketch).

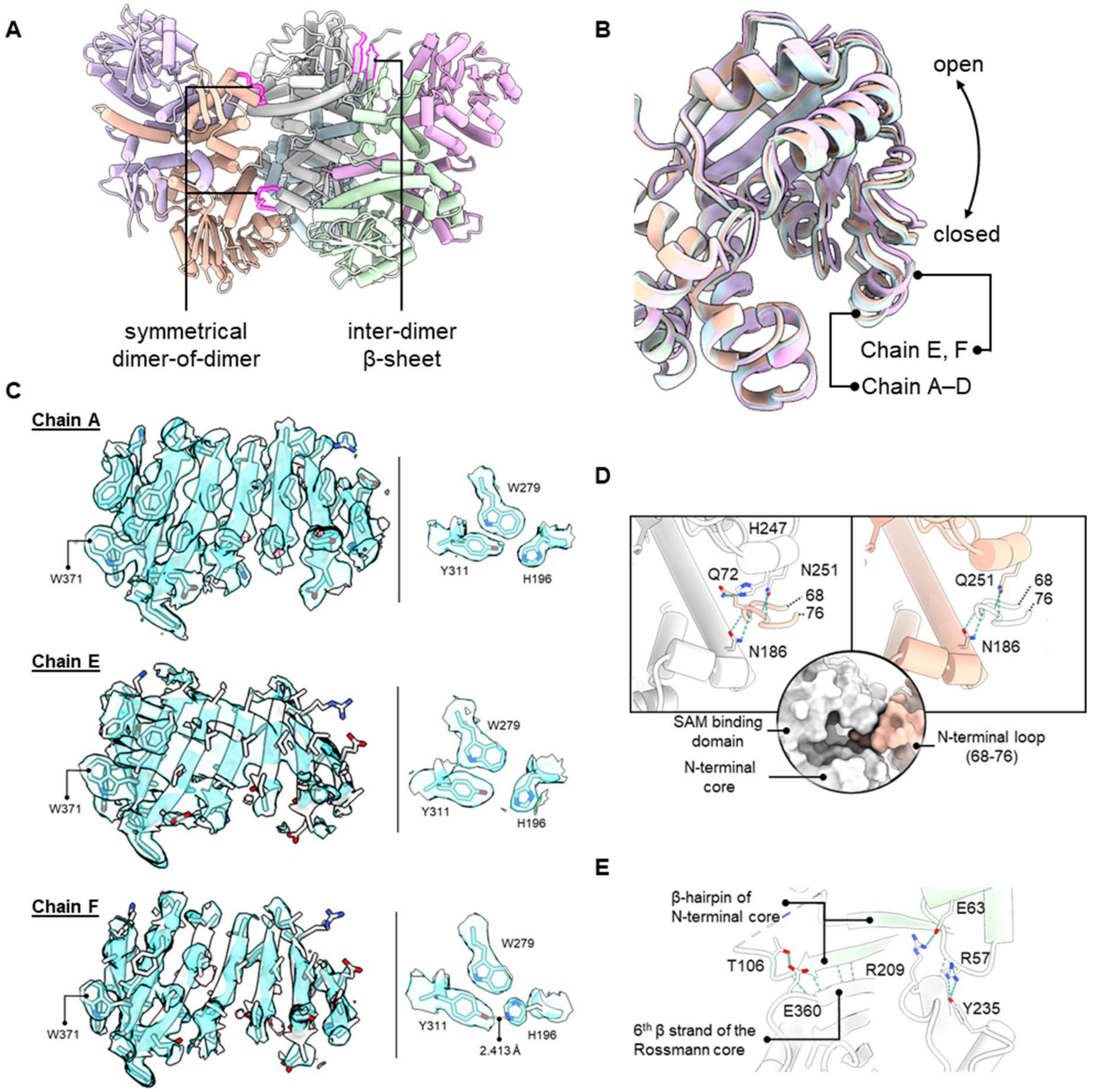


**Fig. S2. Extended structural insights from apo SbSOMT crystal structure.** (**A**) Asymmetrical unit of apo SbSOMT crystal comprises six chains (differentiated by color) forming three pair of dimers. Possible higher order structures are observed (silhouetted in magenta), including the symmetrical dimer-of-dimer and inter-dimer β-sheet. (**B**) Superimposition of all the chains of apo SbSOMT differentiated the openings of SAM binding domain of Chain A–D and of Chain E–F by slight difference. (**C**) 2Fo-Fc maps (blue surface) of Chain A, E, and F focusing on the 7-β-sheet Rossmann fold core (right) and the residues of interest (right, H196, W279 and Y311). The maps are contoured at 1.0σ to highlight the well-resolved density of the residues of interest despite the relatively poor electron density of SAM binding domain of Chain E and F. (**D**) The formation of symmetrical dimer-of-dimer is based on the clamping of SAM binding domain opening towards the opponent dimer's N-terminal loop (68-76), with several hydrogen bonds (blue dotted line) observed. (**E**) The formation of inter-dimer β-sheet is based on the interactions between β-hairpin of the N-terminal core and the 6$^{th}$ β strand of the Rossmann core from two chains.

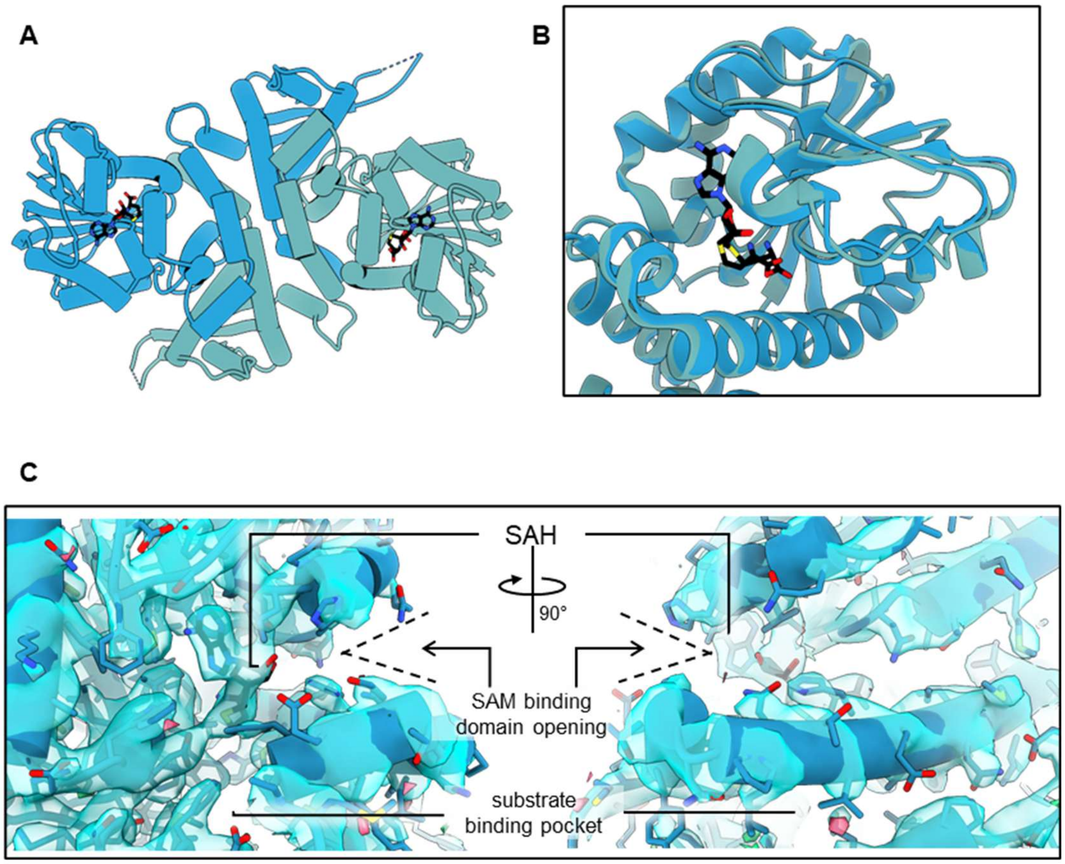


**Fig. S3. Extended structural insights from SAH-bound SbSOMT crystal structure.** (**A**) Asymmetrical unit of apo SbSOMT crystal depicts one dimeric SbSOMT (protomers are differentiated by color). The SAH molecule (black) is bound at the SAM binding pocket of both protomers. (**B**) Superimposition of the protomers highlighting the high similarity of the SAM binding domains and subtle difference of the SAH binding conformations as elaborated in Fig. 3E–F. (**C**) The 2Fo-Fc map (contoured at 1.0σ) at SAM binding domain viewed at two angles; showing the blockage of substrate binding pocket by the bound SAH.

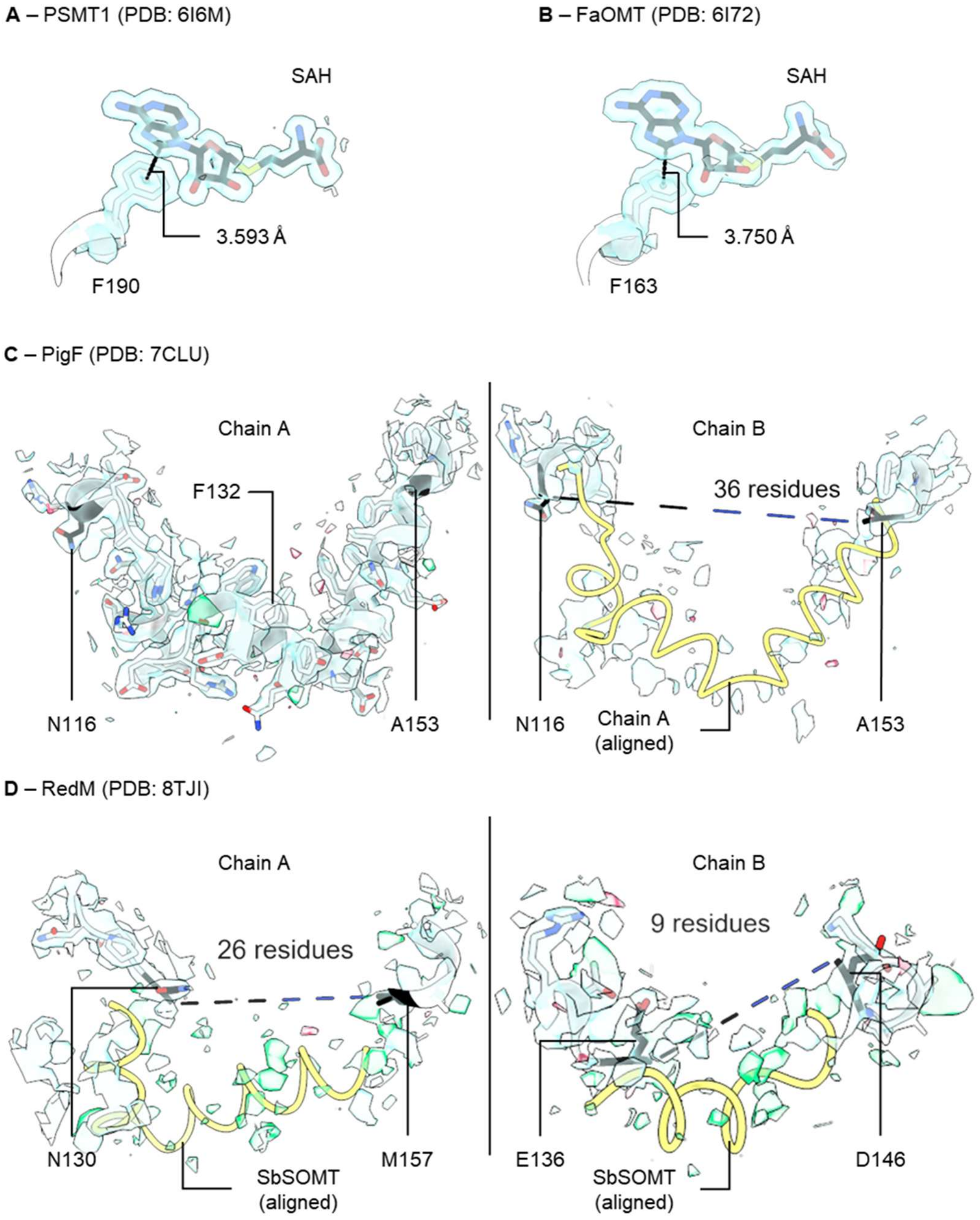


**Fig. S4. Structural insights from SbSOMT homologs on F176-cofactor equivalent π-π interaction and hydrophobic patch instability.** In PSMT1 (**A**) (*21*) and FaOMT (**B**) (*20*),the F176-cofactor equivalent π-π interaction is observed at a shorter distance, 3.593 Å and 3.750 Å, respectively. In Chain B of PigF (**C**) (*22*) and both chains of RedM (**D**) (*23*), the region equivalent to hydrophobic patch of SbSOMT is not modelled, and poor electron density is observed at the surrounding of the unmodelled region. The yellow worm represents the aligned hydrophobic patch region of either Chain A of PigF (**C**) or SbSOMT aligned to RedM (**D**). The unmodelled regions (**C**–**D**) are depicted in dotted lines, and the residues at the ends of the region are labeled and colored in black. The 2Fo-Fc maps (**A**–**D**, blue surface) are contoured at 1.0σ while the Fo-Fc maps (**C**–**D**) are contoured at 3.0σ (green surface) and -3.0σ (red surface).

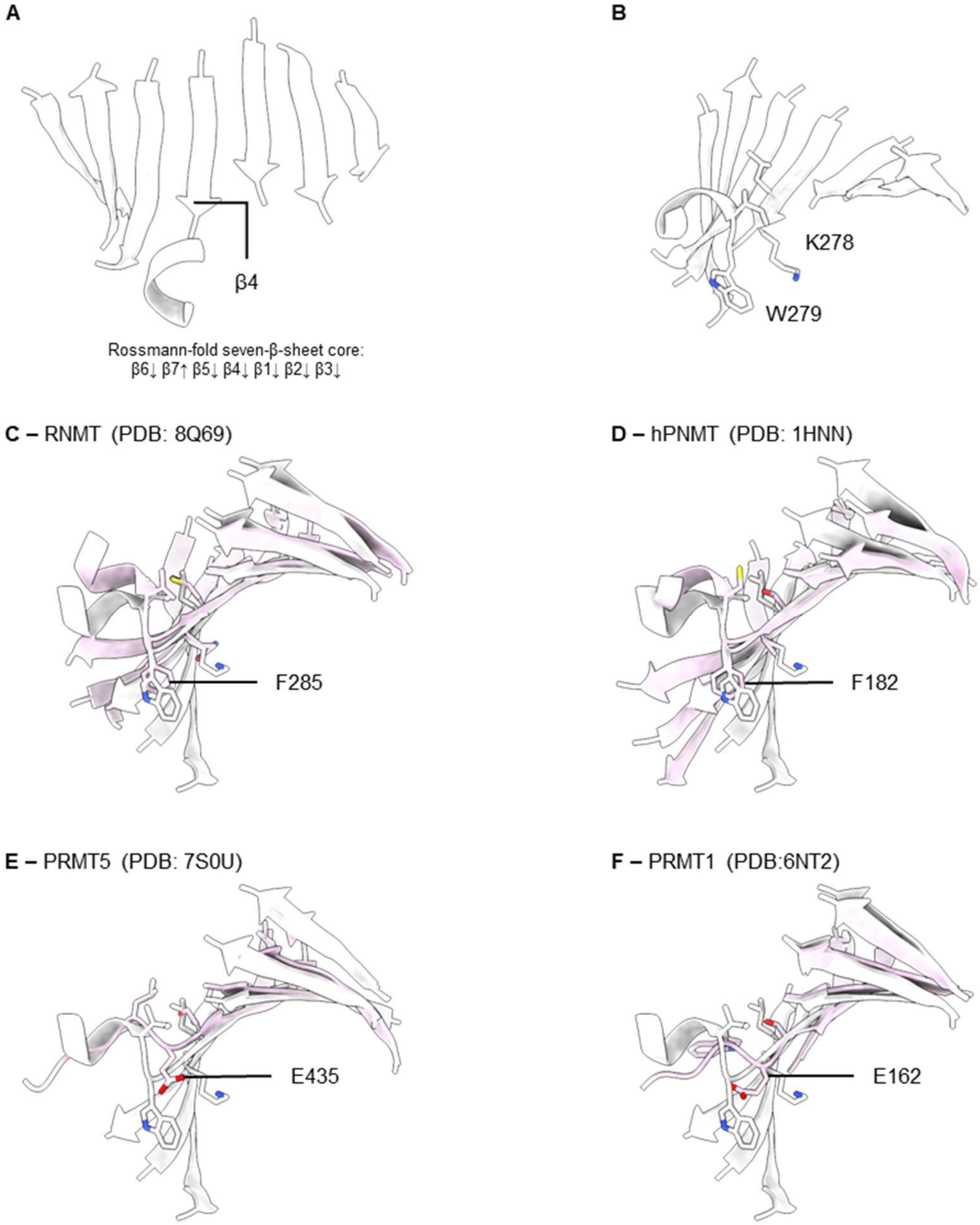


**Fig. S5. Structural comparison of SbSOMT and human Class I homologs.** (**A**) The Rossmann fold seven-β-sheet core of SbSOMT depicting the typical architecture (β6↓β7↑β5↓β4↓β1↓β2↓β3↓) observed in Class I homologs. (**B**) W279 is positioned at the end of β4 strand. W279 is not conserved in RNMT (**C**, F285) (*34*), hPNMT (**D**, F182) (*2*), PRMT5 (**E,** E435) (*33*), and PRMT1 (**F**; E162) (*32*). These human Class I homologs were reported with cooperativity features but lack structural evidence to explain the underlying mechanism.

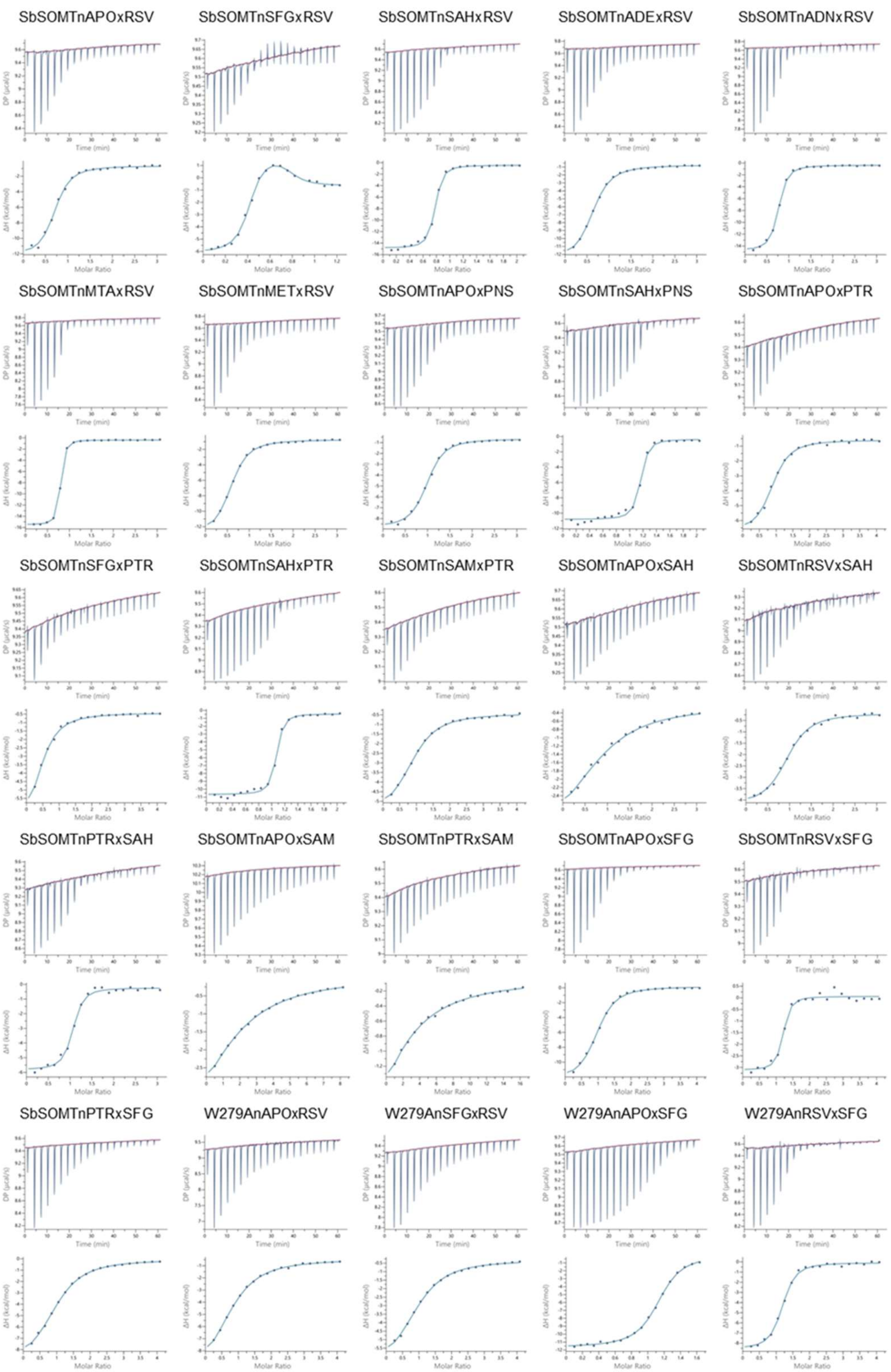


**Fig. S6. Thermograms and integrated plots from ITC.** The label of each set of graphs denotes the sample in cell (SbSOMT or W279A); saturating-ligand in both syringe and cell (n"LIG"); and ligand titrant in syringe (x"LIG"). The red line in the thermogram denotes the baseline set by the software in automated mode for integration of the enthalpy of the injections.

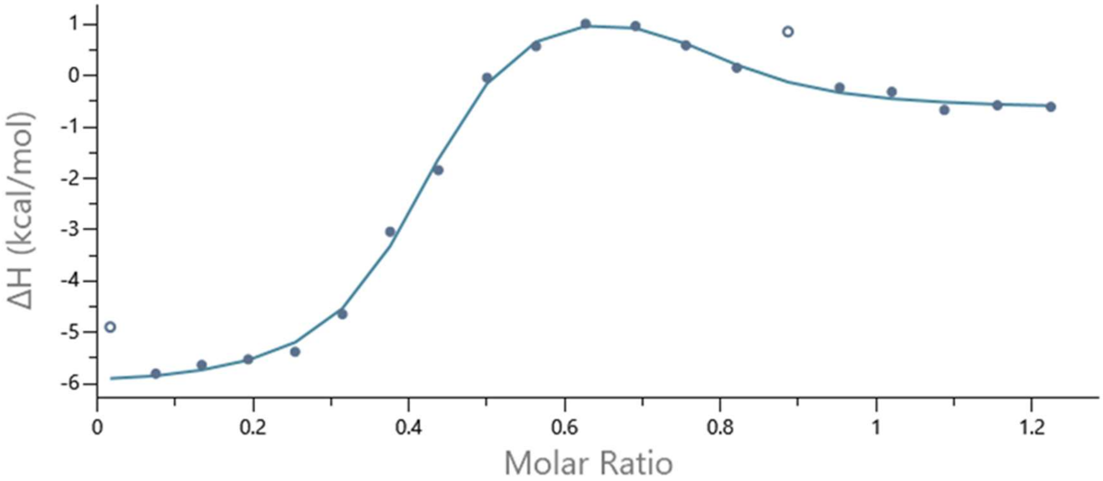


**Fig. S7. ITC integrated plot of RSV binding in SFG-saturated condition.** The plot marks the excluded data points (1st and 15th, open circle) and included data points (closed circle) that were used to integrate the binding kinetics.

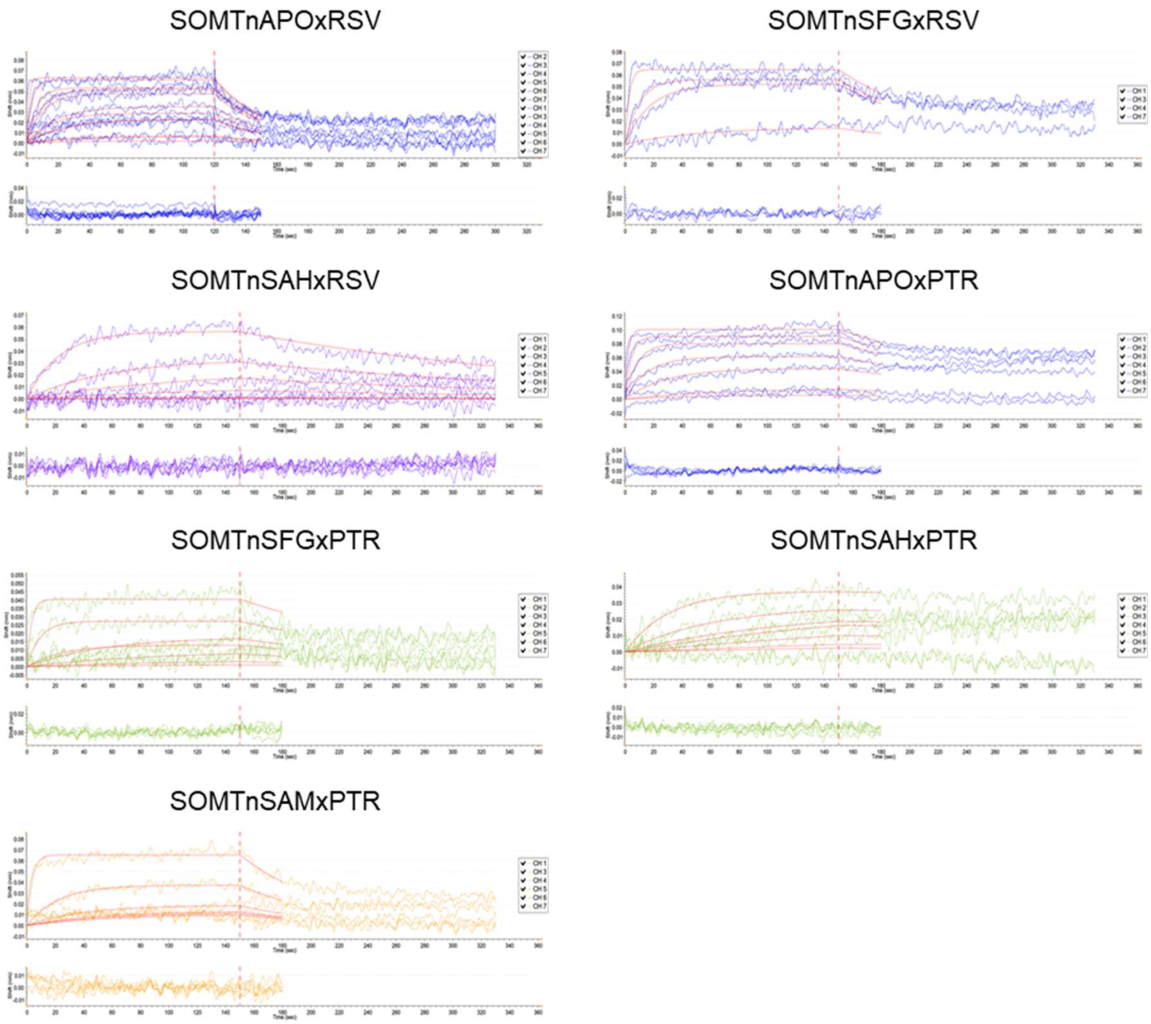


**Fig. S8. BLI sensograms of SbSOMT bindings towards resveratrol and pterostilbene.** Each set of graphs consist binding curve graph (top) and residual curve graph (bottom). The label of each set of graphs denotes the sample in cell (SbSOMT or W279A); saturating-ligand in both syringe and cell (n"LIG"); and ligand titrant in syringe (x"LIG"). The red lines in the binding curve graph are the calculated binding curve fitted globally to the sensograms of selected wells. The red dotted line in both graphs separate the association and dissociation step of the assay.

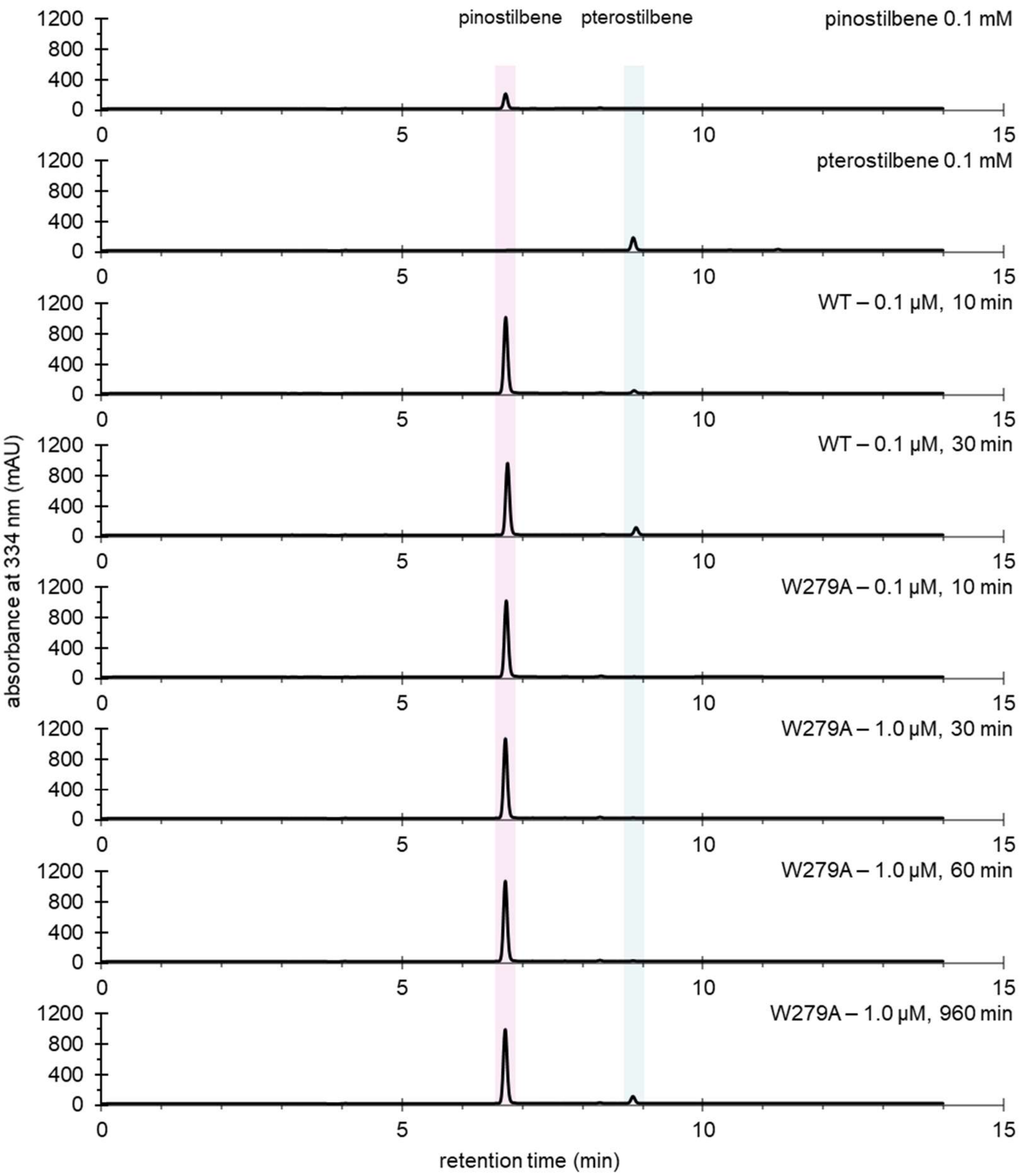


**Fig. S9. Representative HPLC chromatograms of the standards and enzymatic reactions.** The sample identity is labeled at the top right of the chromatogram. The elution peaks of pinostilbene and pterostilbene are highlighted by pink and blue, respectively. WT refers to the wildtype SbSOMT.

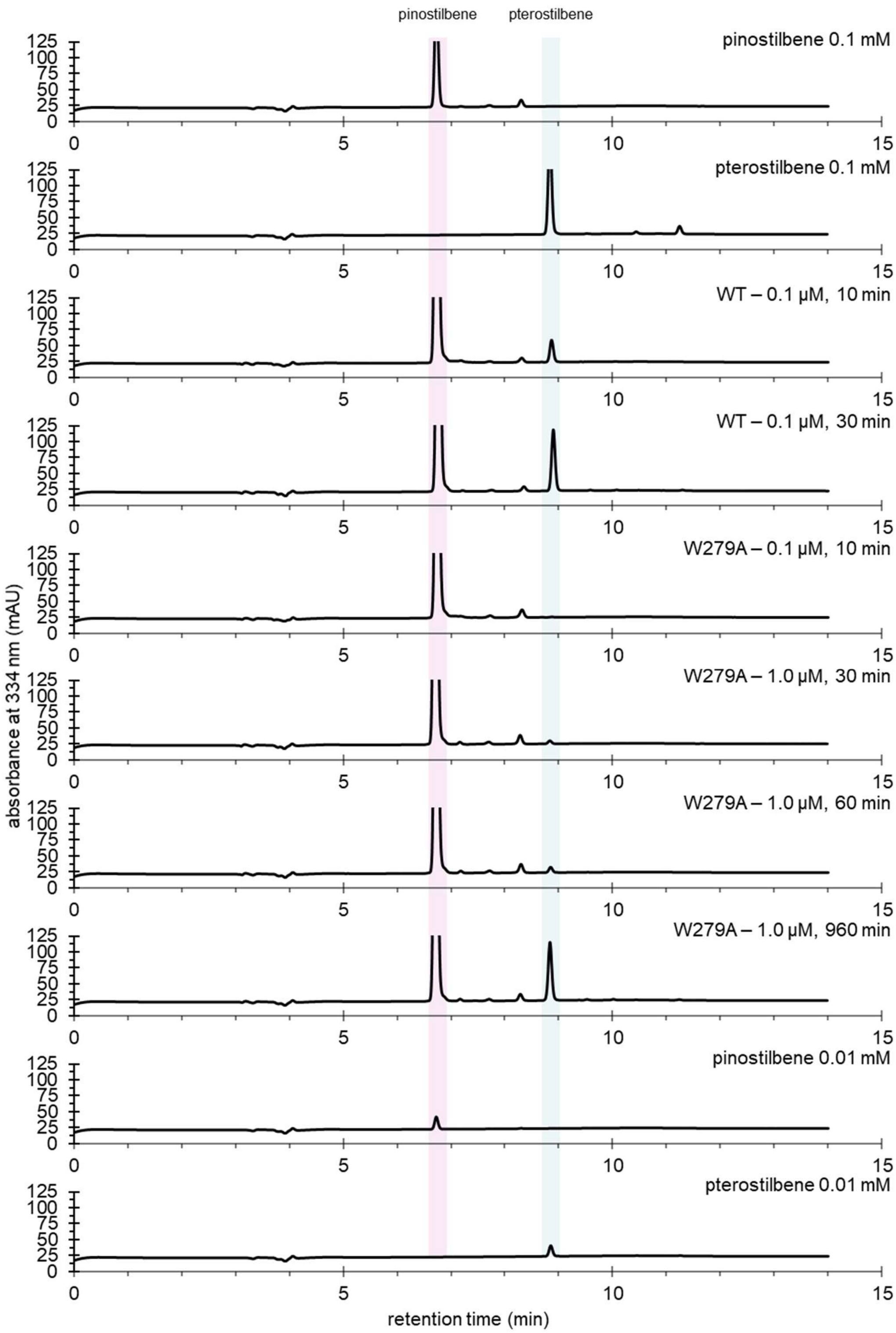


**Fig. S10. Rescaled representative HPLC chromatograms for pterostilbene peak resolution.** The chromatograms are rescaled to resolved the smaller peak of pterostilbene. The sample identity is labeled at the top right of the chromatogram. The elution peak of pinostilbene and pterostilbene are highlighted by pink and blue, respectively. WT refers to the wildtype SbSOMT.

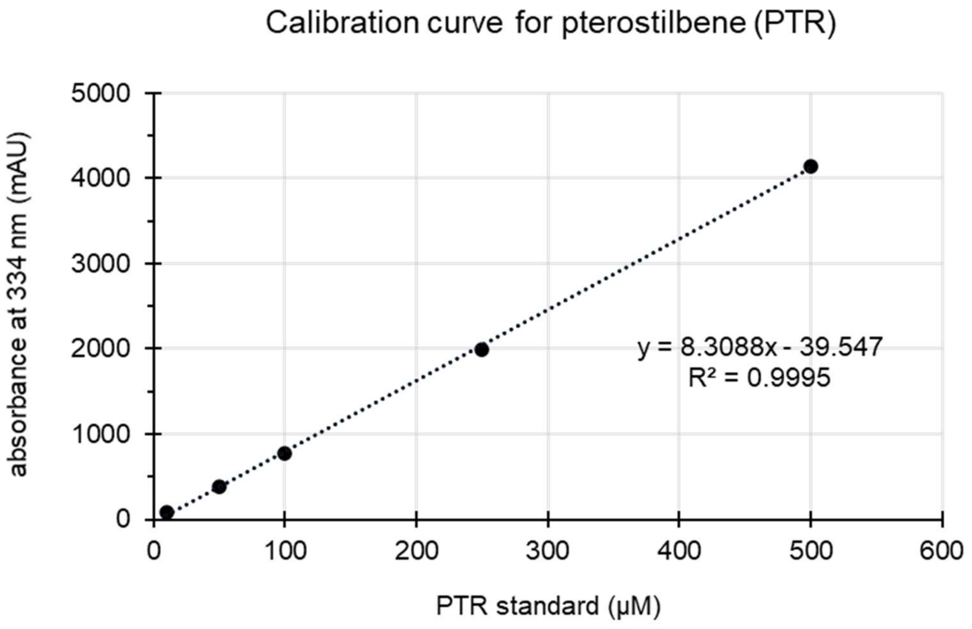


**Fig. S11. Representative calibration curve for quantification of pterostilbene.** Calibration curve for quantification of pterostilbene elution peak area derived from HPLC analysis.

**Table S1. Thermodynamics profile and stoichiometry of W279A derived from ITC.**

| Ligand | Condition[a] | N (sites) | ΔH (kcal/mol) | ΔG (kcal/mol) | -TΔS (kcal/mol) |
|---|---|---|---|---|---|
| RSV | APO | 0.82 ± 0.017 | -10.2 ± 0.406 | -6.12 | 4.12 |
| | SFG+ | 0.97 ± 0.019 | -6.61 ± 0.265 | -6.22 | 0.388 |
| PTR | APO | 1.11 ± 0.006 | -11.6 ± 0.167 | -8.26 | 3.36 |
| | RSV+ | 1.12 ± 0.012 | -8.53 ± 0.155 | -7.88 | 0.653 |

This thermodynamics profile and binding affinity profile at Fig. 4F are derived from the same run and completes each other.

[a]Symbol '+' denotes the ligand is supplied at saturating concentration.

**Table S2. Conditions and setup variables of the ITC runs.**

| Condition | | | Concentration | | | |
|---|---|---|---|---|---|---|
| Cell | Titrant | Constant[a] | Cell (μM] | Titrant (μM) | Constant (μM) | PEG400 (v/v%) |
| SbSOMT | RSV | APO | 750 | 50 | 0 | 2.5 |
| | | SFG | 300 | 50 | 250 | 2.5 |
| | | SAH | 500 | 50 | 250 | 2.5 |
| | | ADE | 750 | 50 | 1000 | 2.5 |
| | | ADN | 750 | 50 | 1000 | 2.5 |
| | | MTA | 750 | 50 | 1000 | 2.5 |
| | | MET | 750 | 50 | 1000 | 2.5 |
| | PNS | APO | 750 | 50 | 0 | 2.5 |
| | | SAH | 500 | 50 | 250 | 2.5 |
| | PTR | APO | 500 | 25 | 00 | 5.0 |
| | | SFG | 500 | 25 | 125 | 5.0 |
| | | SAH | 250 | 25 | 125 | 5.0 |
| | | SAM | 500 | 25 | 500 | 5.0 |
| | SAH | APO | 750 | 50 | 0 | 2.5 |
| | | RSV | 750 | 50 | 250 | 2.5 |
| | | PTR | 750 | 50 | 250 | 5.0 |
| | SFG | APO | 1000 | 50 | 0 | 0 |
| | | RSV | 1000 | 50 | 250 | 2.5 |
| | | PTR | 1000 | 50 | 250 | 5.0 |
| | SAM | APO | 2000 | 50 | 0 | 0 |
| | | PTR | 2000 | 25 | 250 | 5.0 |
| W279A | RSV | APO | 2000 | 100 | 0 | 2.5 |
| | | SFG | 2000 | 100 | 500 | 2.5 |
| | SFG | APO | 400 | 50 | 0 | 2.5 |
| | | RSV | 1000 | 50 | 500 | 2.5 |

[a]Constant refers to the ligand supplied at saturated concentration or 1 mM. In Table 1, Fig. 1, and Fig. 4, these conditions are marked by '+' if the constant is not APO.

**Table S3. Data collection and refinement statistics for crystal structures solved in this study.**

| | Apo SbSOMT | SAH-bound SbSOMT |
|---|---|---|
| **Data collection** | | |
| Wavelength (Å) | 0.979 | 0.979 |
| Resolution (Å) [a] | 80.16 – 2.04 | 59.58 – 2.90 |
| Space group | P 21 21 21 | P 1 21 1 |
| Cell dimensions | | |
| a, b, c (Å) | 84.39 160.07 200.96 | 64.87 94.38 64.99 |
| α, β, γ (°) | 90.00 90.00 90.00 | 90.00 113.71 90.00 |
| Total reflections | 2265827 (95999) | 108296 (18770) |
| Unique reflections | 172235 (8501) | 15690 (2607) |
| Multiplicity | 13.2 (11.3) | 6.9 (7.2) |
| Completeness (%) [a] | 99.3 (100) | 97.7 (100) |
| $I / \sigma I^{a}$ | 6.8(1) | 9.7 (1.6) |
| Wilson B-factor (Å²) | 36.4 | 67.8 |
| $CC_{1/2}$ [a] | 0.995 (0.389) | 0.996 (0.656) |
| **Refinement** | | |
| Reflections used in refinement | 172122 | 15668 |
| Reflections used for R-free | 8845 | 778 |
| $R_{work}$ / $R_{free}$ | 0.217/0.247 | 0.216/0.251 |
| Number of atoms | | |
| Macromolecules | 33310 | 11039 |
| Ligands/Ions | 270 | 88 |
| Solvent | 353 | 12 |
| rmsd [a] bond angles (°) | 0.0129 | 0.0021 |
| rmsd [a] bond lengths (Å) | 2.199 | 0.849 |
| Ramachandran plot | | |
| Favored (%) | 98.71 | 93.64 |
| Allowed (%) | 1.29 | 5.65 |
| Outlier (%) | 0.00 | 0.71 |
| Sidechain outlier (%) | 4.91 | 3.53 |
| Clash score | 5.62 | 2.67 |
| Mean B-value (Å²) | | |
| Macromolecules | 64.33 | 88.55 |
| Ligands | 71.78 | 100.2 |
| Solvent | 51.00 | 52.8 |
| **PDB ID code** | **9WIU** | **9WIV** |

[a] Value relative to the highest resolution shell are given in parentheses.
